# Cortical Sensitivity to Real-World Object Size in Natural Images

**DOI:** 10.1101/2023.03.17.533179

**Authors:** Ahmad Hlayhel, Andrew F. Luo, Leila Wehbe, Michael J. Tarr, Margaret M. Henderson

## Abstract

Real-world size is a fundamental property of objects that supports adaptive interactions with the physical environment. Prior fMRI studies have identified large-scale organization in higher visual cortex associated with real-world object size but have relied primarily on decontextualized objects presented on simple backgrounds. Here, we ask whether this neural organization generalizes to objects embedded within the context of complex natural scene images. To address this question, we used voxelwise analyses and encoding models to assess neural responses to natural images from two independent fMRI datasets. These analyses identified reliable organization encompassing alternating regions of sensitivity for small and large objects across the ventral and lateral surfaces of occipitotemporal cortex, which largely aligns with prior findings using isolated objects. At the same time, we identify additional, previously unreported size-related organization on the ventral surface at a finer spatial scale. We also find real-world size sensitivity for both animate and inanimate objects, with broadly corresponding cortical patterns across the two domains. Importantly, controlling for low- and mid-level visual features, object composition, and scene clutter attenuated the magnitude of size sensitivity in some regions, but closely preserved the spatial organization and directionality of neural size preferences. Finally, we characterized the shape of voxelwise tuning functions for real-world object size, and found that a logarithmic-like relationship predominated, although profile shape varied across regions and between small- and large-preferring voxels. Overall, our findings support real-world object size as a behaviorally-relevant dimension of cortical organization during natural vision.

## 1 Introduction

The human visual system represents the objects and scenes around us in ways that support adaptive behavior. This functional organization is reflected in spatially localized regions of higher visual cortex that are selective for particular categories such as faces, words, scenes, tools, and food (Cohen et al., 2000; R. Epstein and Kanwisher, 1998; Jain et al., 2023; Kanwisher et al., 1997; Mahon et al., 2007; Sergent et al., 1992) as well as large-scale organizing gradients based on object properties such as object animacy, real-world size, and reachability (Gallivan et al., 2009; Josephs & Konkle, 2020; Konkle & Caramazza, 2013; Naselaris et al., 2012; Warrington & Shallice, 1984). Real-world object size – how large an object is inferred to be in the physical world, as opposed to its size in the retinal projection – plays an important role in interacting with objects (e.g., mistaking tigers for house cats rarely ends well).

Mechanisms of size constancy stably encode real-world object size across variation in properties like viewpoint and distance Sperandio and Chouinard (2015), allowing us to make inferences about the affordances of objects. For example, we perceive small objects as things we can pick up and larger objects as things to be navigated around (Gibson, 1950; Grèzes et al., 2003; Slater et al., 1990; Tucker & Ellis, 2001).

Despite the clear behavioral importance of object size, studies demonstrating neural sensitivity to real-world size (e.g., Blumenthal et al., 2018; Julian et al., 2017; Konkle & Caramazza, 2013; Konkle & Oliva, 2012) have typically relied on highly controlled experimental designs in which objects are presented in isolation, without the naturalistic backgrounds or scene context that provide visual and contextual cues to object size. In natural vision, by contrast, judgments of real-world size are influenced by background information and other objects in the scene (Bar, 2004). For example, the 3D Ponzo illusion shows that objects with the same retinal size, but presented in different depth contexts, can be perceived and cortically represented differently (Murray et al., 2006). Given the apparent importance of context for inferring real-world size, we explore whether cortical sensitivity to this property – previously characterized primarily using isolated objects – is also evident in complex natural scenes.

To address this question, we first tested whether single-voxel cortical responses exhibit sensitivity to real-world object size in complex natural scenes using two large-scale, naturalistic fMRI datasets, the Natural Scenes Dataset (NSD) and BOLD5000 (Allen et al., 2022; Chang et al., 2019). To establish the real-world size category for each natural image, we used a computational saliency model (Jiang et al., 2015) to identify the primary object in the image and then assigned the image into the appropriate size category (Large or Small; Fig. 1). We then characterized the topographic cortical organization of real-world size sensitivity and compared our estimates to the spatial patterns reported in past work.

**Figure 1:**
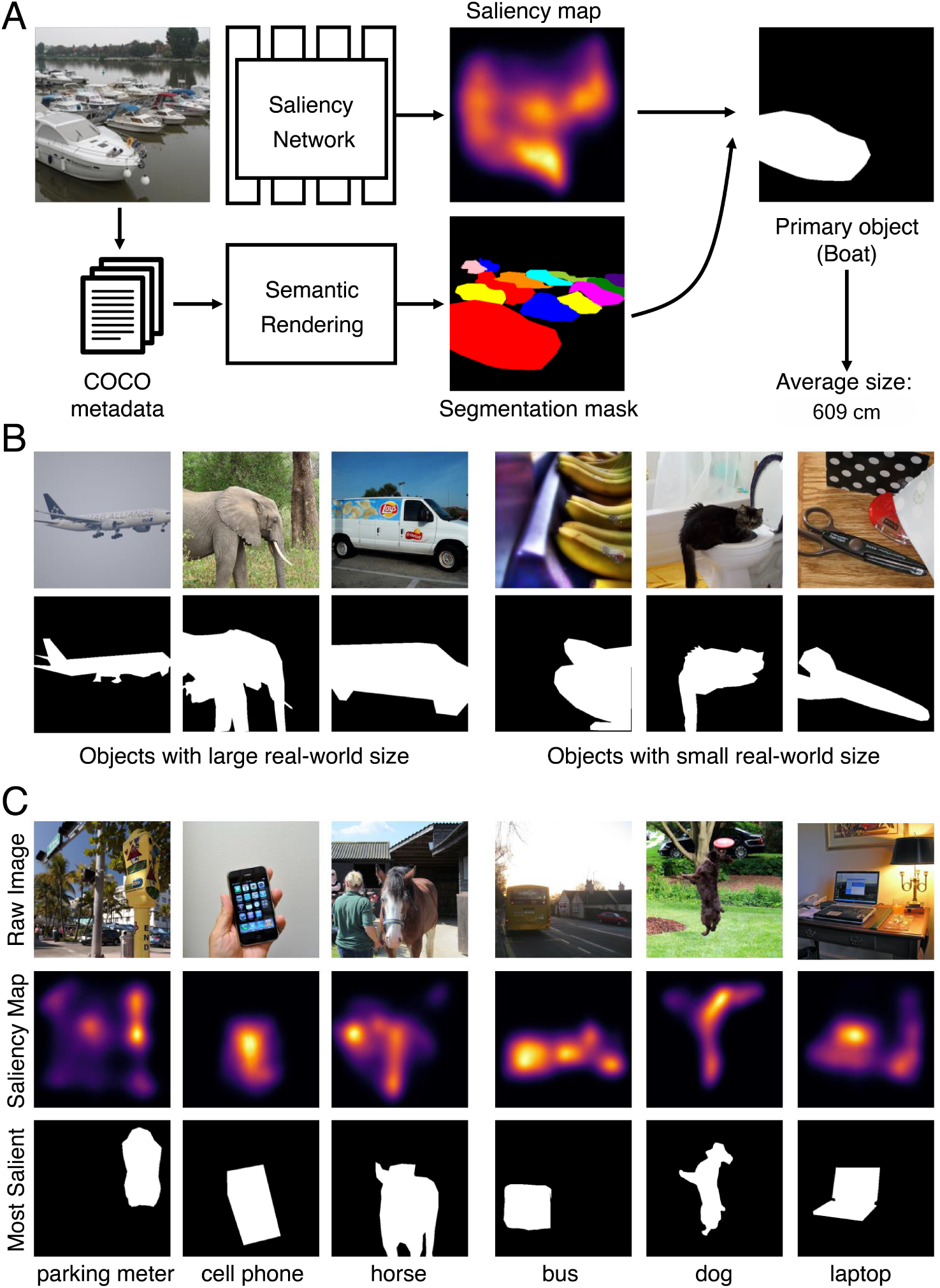
Labeling real-world size in natural scenes using saliency-guided primary object identification. **(A)** An image is passed to a convolutional neural network trained to predict human saliency from the SALICON dataset (Jiang et al., 2015). By combining object instance masks (Lin et al., 2014) with the predicted saliency map, we identify the most salient object in each image and label the image based on the estimated real-world size of that object category (see *Methods* for details). **(B)** Examples of objects with large and small real-world size. Original color stimuli are shown on top, with the corresponding binary object masks shown below. **(C)** Example outputs of our saliency extraction pipeline.

Second, motivated by a prior study which reported size sensitivity for inanimate objects but not animate objects (Konkle & Caramazza, 2013), we compared real-world size sensitivity separately for animate and inanimate objects embedded in natural scenes.

Third, we asked whether there is cortical sensitivity to real-world size after accounting for other visual properties associated with object size. Specifically, large and small objects can differ in lower- and mid-level properties, including curvature and boxiness (Nasr & Tootell, 2012; Nasr et al., 2014) as well as orientation and spatial frequency (Henderson et al., 2023). Moreover, some past work suggests that mid-level textural features may be able to account for behavioral judgments of object real-world size (Long & Konkle, 2017; Long et al., 2016), and that images containing only mid-level features can elicit patterns resembling real-world size sensitivity for intact objects (Long et al., 2018). Natural scenes also introduce covariates that may contribute to neural responses, including background context, object area, clutter, and the presence of multiple objects. To address these potential confounds, we evaluated size sensitivity while controlling for several low- and mid-level feature spaces, for alternative rules for defining the primary object, and for object-layout covariates. We further compared encoding models based on the visual features alone with encoding models that additionally included real-world size, asking whether real-world size increased the held-out variance explained in voxel responses. Fourth, we characterized the relationship between real-world size and cortical responses and asked whether its shape differs between small- and large-preferring voxels. Past studies have primarily identified which regions exhibit qualitative differences in real-world size sensitivity (i.e., preference for large or small) (Julian et al., 2017; Konkle & Caramazza, 2013; Konkle & Oliva, 2012). Because real-world object sizes span several orders of magnitude, characterizing the shape of this relationship can reveal whether small- and large-preferring populations differ not only in preference direction, but also in which portions of the size range they differentiate most strongly. Here, we quantitatively extended this approach by comparing the relative held-out fits of logarithmic-like, linear, and exponential-like monotonic functions and testing whether the distribution of best-fitting functions differs between small- and large-preferring voxels. Previewing our findings, voxelwise analyses using natural images reveal a pattern of real-world size sensitivity that corresponds to cortical patterns reported in past work, with some differences in the fine-grained organization. We also observed reliable sensitivity to real-world size in several category-selective cortical regions. In contrast to prior work, we find that size sensitivity exhibited a similar spatial organization for animate and inanimate objects. Importantly, the overall pattern of real-world size sensitivity remains detectable after accounting for low- and mid-level visual feature spaces, alternative object-definition rules and object-layout covariates, suggesting that these potential confounds do not fully account for the observed real-world size sensitivity. Finally, a voxelwise comparison of three monotonic neural response functions (logarithmic-like, linear, and exponential-like) revealed that size tuning profiles differ between small- and large-preferring voxels and across cortical regions.

## 2 Materials & Methods

### fMRI data

We used the Natural Scenes Dataset (NSD; Allen et al. (2022)). Additional visualization and analyses used BOLD5000 (Chang et al., 2019). The detailed data collection procedures can be found in Allen et al. (2022) and Chang et al. (2019). In the NSD, human participants viewed natural images sourced from the Microsoft Common Objects in Context (MS COCO) dataset (Lin et al., 2014), which include instance segmentation annotations. Whole-brain fMRI data were collected using a 7T scanner. Eight subjects each viewed between 9,209 and 10,000 images, with each image repeated approximately three times. Of 70,566 total unique images presented, a total of 907 images were viewed at least once by all subjects. Single-trial betas were generated using GLMsingle (Prince et al., 2022). This process estimated a per-voxel hemodynamic response function. Cross-validation and ridge regression steps were further applied to estimate the beta coefficients. We computed the mean and standard deviation of each voxel across a given scanning session and used these values to compute a *z*-value for each voxel corresponding to each trial. *Z* -values were averaged across repetitions of an image (approximately 3 trials/image) to compute an averaged response value for each voxel/image combination. In several analyses, we summarized our data using functionally defined regions of interest (ROIs). These included early visual areas (V1, V2, V3, hV4), scene-selective areas (the retrosplenial cortex [RSC], parahippocampal place area [PPA], and occipital place area [OPA]), body-selective areas (the extrastriate body area [EBA] and fusiform body area [FBA]), face-selective areas (the occipital face area [OFA] and fusiform face area [FFA]), and word-selective areas (the occipital word form area [OWFA] and visual word form area [VWFA]). These ROI definitions were obtained using independent retinotopic mapping and category localizer scans performed as part of the NSD experiment (see Allen et al. (2022) for details on ROI definitions).

BOLD5000 data were collected from four subjects using a 3T scanner, with stimuli drawn from three image sources: MS COCO (Lin et al., 2014), ImageNet (Russakovsky et al., 2015), and SUN (Xiao et al., 2010). For the primary cross-dataset comparison, we restricted our analyses to MS COCO stimuli, allowing us to apply the same COCO-based image-labeling procedure used for NSD. In a supplementary analysis, we also analyzed the ImageNet stimuli to assess whether the observed size-sensitivity organization extended beyond MS COCO (see Supplementary Material). We estimated single-trial beta coefficients using GLMsingle and applied the same normalization and z-scoring procedure used for NSD. Both NSD and BOLD5000 are publicly available datasets; participant recruitment, informed consent, and ethics approval for the original data collection are described in the corresponding source publications (Allen et al., 2022; Chang et al., 2019). The present work is a secondary analysis of de-identified, publicly available data and did not involve new data collection.

### Real-world object size assignment

MS COCO provides instance segmentation annotations for object instances belonging to 80 semantic categories. As the first step in assigning a real-world size label to each image, we assigned each semantic object category in MS COCO a real-world size estimate. To obtain these estimates, we queried a large language model (OpenAI GPT-4o) for the physical dimensions of each of the 80 MS COCO object categories. For each category, we collected ten independent responses, each reporting an object’s length, width, and height in centimeters. From each response, we retained the largest physical dimension and averaged this value across the ten responses to obtain a single category-level size estimate. The exact prompt is provided in the Supplementary Material. We then reviewed the resulting estimates, flagging categories with high variability across the ten responses and manually correcting a small number of clear errors. These estimates represent the typical real-world size of each semantic category rather than a measurement of the specific exemplar depicted in an image.

### Saliency-guided object selection

Many of the images in our dataset include multiple objects within a single image. To allow each image to be assigned a single real-world size label, we selected the single most salient object within each image. Visual saliency was estimated using SimpleNet, a PNASNet-5-based encoder–decoder network trained on the SALICON dataset (Jiang et al., 2015; Reddy et al., 2020). SALICON provides human attention maps derived from mouse-tracking during free viewing of MS COCO images. The saliency prediction was represented as a single-channel image with values in [0, 1] and spatially aligned with the input image. Individual object masks were rendered from MS COCO metadata. For each object, we computed a spatial integral of the saliency over the area defined by its mask. The object with the maximum saliency integral was retained. This primary-object designation does not imply that neural responses are driven only by the selected object. We therefore assessed the robustness of the results using alternative object-definition rules and controls for primary-object area, the number of objects, and non-primary-object area (see below). To avoid degenerate objects, we excluded images in which the most salient object’s mask occupied less than 0.5% of the image area. Our approach is illustrated in Fig. 1.

The final real-world size label for each image was then determined using the estimated real-world size (see previous paragraph) of the object category corresponding to the most salient object. In all analyses, we followed Konkle and Oliva (2012) and excluded any images in which the most salient object was a person in order to avoid potential biasing effects of this label (“person” is the most common object label in MS COCO by a substantial margin, and images of people are known to evoke strong cortical responses in multiple brain areas; e.g., Kanwisher et al. (1997)).

Our combined criteria yielded between 5, 944 and 6, 583 images per subject in NSD, and between 796 and 1, 328 MS COCO images per subject in BOLD5000. In our analysis of animate versus inanimate objects, we defined an object class as animate if it was an animal. For example, objects such as Cat and Dog were treated as animate, while Apple and Car were treated as inanimate. Animacy assignment for an image was similarly based on the most salient object.

Across subjects, the final NSD image sets contained between 4, 369 and 4, 847 inanimate images and between 1, 575 and 1, 737 animate images.

### Size sensitivity analysis

To identify size-sensitive voxels, we used a cross-validated procedure conducted separately for each subject. We first divided each subject’s valid images into small- and large-object groups using the median real-world size (169 cm) and generated ten random split-half partitions, where each partition had comparable numbers from the large- and small-object groups in both halves. Within each partition, one half served as the training set for voxel selection and the other as the held-out set for estimating size sensitivity, after which their roles were reversed. For voxel selection, the median was recomputed using only the training images, which were then divided into small- and large-object groups. For each voxel, we compared the mean response between these groups using a two-sample, two-tailed *t*-test. The resulting *p*-values were corrected across all cortical voxels within each subject and fold using the Benjamini–Hochberg false discovery rate (FDR) procedure (Benjamini & Hochberg, 1995) (*q <* 0.05). Across the ten partitions and two fold directions, this procedure yielded 20 selection–estimation folds. A voxel was considered reliably size-sensitive if it met the training-set selection criterion in more than half of the folds (i.e., at least 11 of 20 folds).

On each held-out half, we estimated size sensitivity using Spearman’s rank correlation between each voxel’s response and the continuous real-world size estimate assigned to the primary object. For each voxel, correlations were averaged across the 20 held-out folds. For ROI-level analyses, we averaged the held-out correlations across all voxels within each independently localized ROI. The resulting subject-level ROI mean sensitivity values were compared against a null hypothesis of zero using two-tailed one-sample *t*-tests across the eight subjects, with Benjamini–Hochberg FDR correction (*q <* 0.05) across ten ROIs: pooled V1–hV4, RSC, PPA, OPA, EBA, FBA, OFA, FFA, OWFA, and VWFA. For visualization, we projected values onto cortical surface meshes using PyCortex software (Gao et al., 2015). For group-level visualization, cortical data were projected onto a common fsaverage surface.

### Controlling for low- and mid-level visual features

In order to address possible contributions of low- and mid-level visual features to real-world size sensitivity, we constructed voxel-wise encoding models (Naselaris et al., 2011) based on several low- and mid-level feature spaces: a Gabor model (Henderson et al., 2023), a GIST model (Oliva & Torralba, 2001), and a “sketch tokens” model (Lim et al., 2013). Both the Gabor and GIST models were used as examples of low-level feature spaces, as they are each based on orientation and spatial frequency features, while the sketch tokens model was used as a mid-level feature space, as it utilizes a model that learns to categorize image contour patches into a set of data-driven clusters (Lim et al., 2013). In addition to these three models, we constructed a combined model by concatenating the Gabor, GIST, and sketch tokens feature spaces, yielding a 759-dimensional feature vector. For both the Gabor and sketch tokens models, we used a fitting procedure that takes into account the spatial receptive field of each voxel, or its population receptive field (pRF). An estimate of each voxel’s pRF was obtained based on an independent retinotopic mapping task performed as part of the NSD experiment (see Allen et al. (2022) and Benson et al. (2018) for details on the task and pRF estimation). The parameters of this estimated pRF (a 2D Gaussian with parameters (*µ_x_, µ_y_*) = center, *σ* = size) were then quantized by rounding the original parameter value to the nearest value in a pre-specified grid over candidate pRF values (this quantization was performed to reduce computing time). Within the pRF parameter grid, the size and eccentricity values were each logarithmically distributed with 10 possible values each, with eccentricity ranging from 0*^◦^* to 7*^◦^* (*^◦^* = degrees visual angle), and size ranging from 0.17*^◦^* to 8.4*^◦^*. Angular position values were evenly spaced along 16 possible angular values. The quantized parameterization yielded a total of 1, 456 possible size/eccentricity/angular position combinations (combinations whose receptive fields did not overlap the image extent were excluded). The pRF estimates were then used to compute a single feature vector for each voxel and each image, by taking a weighted sum between the voxel’s pRF (a 2D Gaussian) and a stack of feature maps corresponding to the image of interest (see St-Yves and Naselaris (2018) and Henderson et al. (2023) for details on this approach). For the Gabor model, these maps were computed at 12 orientations and 8 spatial frequencies, for a total of 96 spatial feature maps (see Henderson et al. (2023) for more details on model construction). For the sketch tokens model, there were 151 spatial feature maps, each indicating the presence of a specific class of contour feature. For the GIST model, we did not include an estimate of the voxel’s pRF, because this model incorporates features from the entire image. The GIST model was implemented following the procedure described by Oliva and Torralba (2001); we used an implementation with a 4 *×* 4 spatial grid, 4 spatial frequencies, and 8 orientations, for a combined 512-dimensional feature vector.

For each visual feature model, we tested whether the association between voxel responses and real-world size remained after accounting for the corresponding feature representation by correlating their held-out residuals after regressing the visual features from both quantities. Using the split-half partition described above, we fitted two separate linear models on the training images to predict (i) the real-world size label of the image and (ii) each voxel’s response based on the model’s visual features. We then applied these models to the held-out images and computed residuals for each quantity as the observed value minus the feature-predicted value. Post-control size sensitivity was quantified as the Spearman rank correlation between the held-out residual size values and residual voxel responses. Each image half served once as the training set and once as the held-out set, and the two resulting correlation estimates were averaged. For direct comparison, raw size sensitivity was computed based on the correlation between real-world size and the original voxel responses on the same held-out images, without residualization. We additionally computed the held-out variance explained (*R*^2^) by each feature model to quantify how well the corresponding feature representation predicted voxel responses. For ROI-level analyses, each subject’s post-control correlations were averaged across all voxels within each independently defined ROI. Group-level significance was assessed using two-tailed one-sample *t*-tests across the eight subjects, with Benjamini–Hochberg FDR correction (*q <* 0.05) across the 52 post-control tests comprising four feature models and 13 ROIs: V1, V2, V3, hV4, RSC, PPA, OPA, EBA, FBA, OFA, FFA, OWFA, and VWFA.

The raw baseline shown for comparison was evaluated separately using the same two-tailed one-sample *t*-tests, with Benjamini–Hochberg FDR correction (*q <* 0.05) across the 13 ROIs, and was not included in the 52-test feature-control family. For cortical visualization of post-control size sensitivity, we used the reliable-voxel criterion from the original size-sensitivity analysis rather than reselecting voxels separately after feature control. As a complementary analysis, we tested whether adding real-world size improved held-out prediction of voxel responses beyond the tested visual features. For each voxel and image, Gabor, sketch token, and GIST features were concatenated into a combined visual feature representation. We fitted one encoding model using these visual features alone and a second using the same features together with real-world size. Both models were fitted on the training images and evaluated on the held-out images using the same split-half partition described above. For each voxel, the incremental predictive contribution of real-world size was quantified as 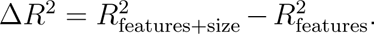 Subject-level Δ*R*^2^ values were averaged across all voxels within each independently defined ROI, without preselecting voxels based on size sensitivity. Group-level significance was assessed using two-tailed one-sample *t*-tests across the eight subjects, with Benjamini–Hochberg FDR correction (*q <* 0.05) across the 13 tested ROIs.

### Controlling for object selection and object layout

Because each image was summarized by the single most salient object in the primary analysis, we tested whether the results were robust to alternative object-definition rules and to properties of the other objects in the scene. We recomputed the image-level real-world size label using three alternative rules in addition to the saliency-primary definition: (i) a largest-object label, defined by the valid object with the largest mask area; (ii) an area-weighted label, defined as the mask-area-weighted average of the log real-world sizes of all valid objects in the image; and (iii) a saliency-weighted label, defined as the corresponding average weighted by each object’s saliency integral (Fig. 7). For each object-definition rule, we computed the voxelwise Spearman rank correlation between real-world size and voxel response across all images that yielded a valid label under the criteria described above. Correlations were then averaged across all voxels within each independently defined ROI. Subject-level ROI values were evaluated using two-tailed one-sample *t*-tests across the eight subjects, with Benjamini–Hochberg FDR correction (*q <* 0.05) across the 52 tests comprising four object-definition rules and 13 ROIs. To assess potential contributions of object layout and scene clutter, we used four control models containing regressors for: (i) the fraction of the image occupied by the primary object’s mask; (ii) the number of usable object masks; (iii) the fraction of the image occupied by non-primary object masks; and (iv) all three variables jointly. Using the same held-out residualization procedure as for the visual-feature controls, we fitted separate linear models on the training images to predict real-world size and each voxel’s responses from the layout variables. The fitted models were applied to the held-out images, and residuals were calculated as the observed values minus the corresponding predictions. Post-control size sensitivity was quantified as the Spearman rank correlation between the held-out size residuals and voxel-response residuals. Each image half served once as the training set and once as the held-out set, and the two resulting correlation estimates were averaged. For ROI-level analyses, each subject’s post-control correlations were averaged across all voxels within each independently defined ROI. Group-level significance was assessed using two-tailed one-sample *t*-tests across the eight subjects, with a separate Benjamini–Hochberg FDR correction (*q <* 0.05) across the 52 tests comprising four layout-control models and the same 13 ROIs. The raw baseline shown for comparison was computed using the same held-out images and fold averaging as the feature-control baseline, evaluated separately with Benjamini–Hochberg FDR correction (*q <* 0.05) across the 13 ROIs, and was not included in the 52-test layout-control family.

### Size tuning profiles

To characterize the shape of the relationship between real-world size and neural response, we compared a set of candidate monotonic transforms of real-world size. Physical size in centimeters was normalized to the unit interval *x ∈* [0, 1] using the minimum and maximum across each subject’s valid images, and we evaluated functions of the form *f* (*x*) = *x^γ^* with *γ ∈ {*1*/*10, 1*/*5, 1*/*2, 1, 2, 5, 10*}*.

Values of *γ <* 1, *γ* = 1, and *γ >* 1 correspond to logarithmic-like, linear, and exponential-like transforms, respectively. For each voxel and candidate *γ*, we fitted a single-predictor linear model (*β ≈ a* + *b x^γ^*) on the training images and evaluated its coefficient of determination (*R*^2^) on the held-out images. The candidate yielding the highest held-out *R*^2^ was retained for that voxel and fold.

For each subject, ROI, and preference group, we computed the fraction of selected voxels assigned to each function class separately within each fold and then averaged these fractions across the ten split-half partitions and both fold directions. We tested whether logarithmic-like transforms constituted more than half of the selected voxels within each ROI and preference group using one-sided one-sample *t*-tests against a fraction of 0.5, with Benjamini–Hochberg FDR correction (*q <* 0.05) across the 20 ROI-by-preference comparisons. We separately tested whether the fraction of exponential-like fits differed between small- and large-preferring voxels using paired two-tailed *t*-tests across the eight subjects, with Benjamini–Hochberg FDR correction (*q <* 0.05) across the ten ROIs.

## 3 Results

### 3.1 Analysis 1: Cortical sensitivity to real-world object size in natural images

To characterize the topographic organization of size-sensitive cortical responses, we first examined the size sensitivity of voxels across higher visual cortex using the Natural Scenes Dataset (Allen et al., 2022). Real-world size labels were assigned to each NSD image by computationally estimating the physical size of the single most salient object per image (see *Methods* and Figure 1 for more details). We identified size-sensitive voxels using a cross-validated procedure in which voxels having significant size sensitivity were identified based on a differential response to large and small objects (two-sample two-tailed *t*-test), and the direction and magnitude of size sensitivity were then estimated for each significant voxel on held-out images using the Spearman rank correlation between voxel response and object size (see *Methods* for more details). Group-level maps are presented in Fig. 2A, while individual-subject results are presented in Fig. 3.

**Figure 2:**
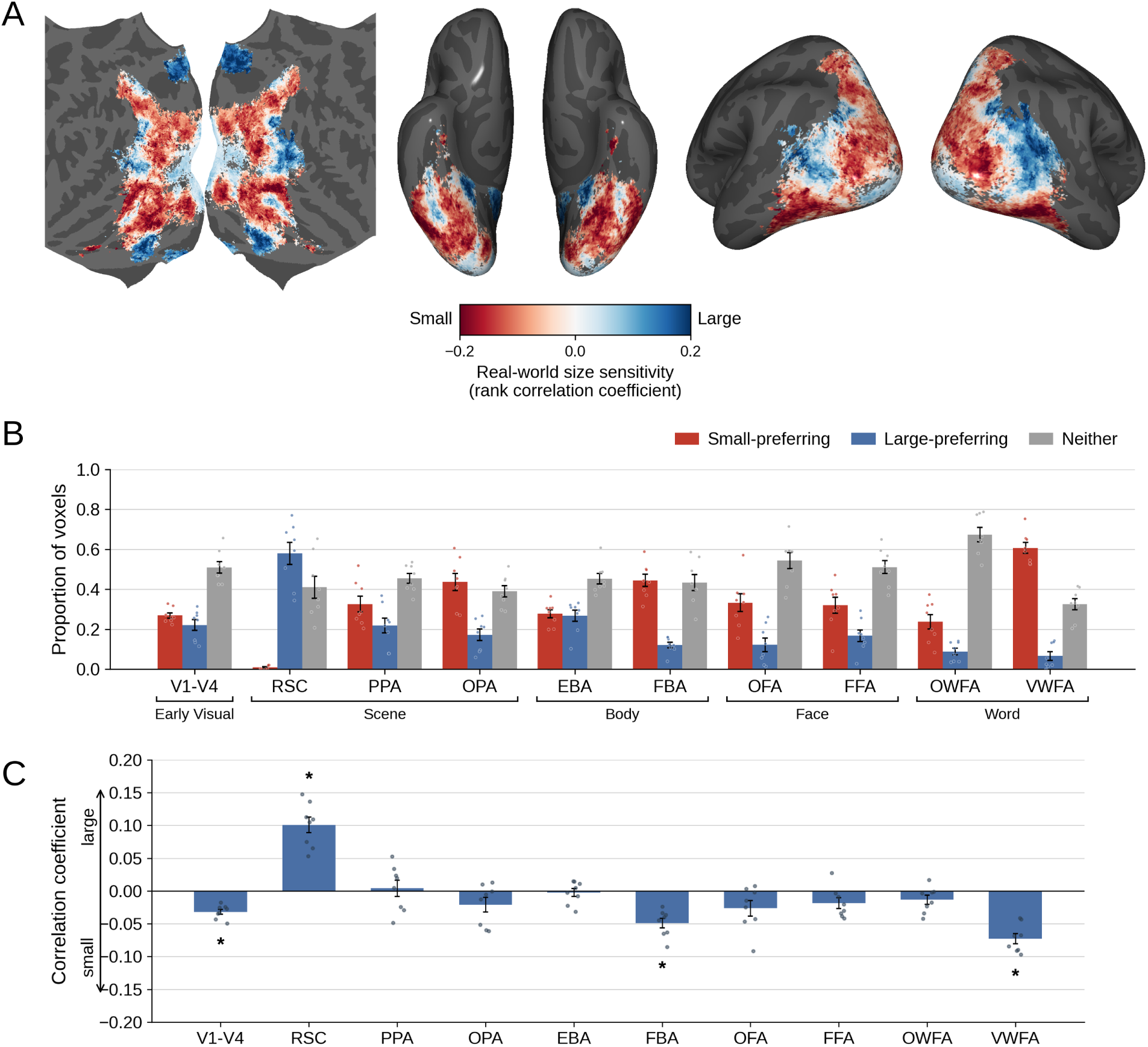
Cortical sensitivity to real-world object size. **(A)** Group size-sensitivity maps in NSD, shown in flattened, ventral, and inflated views. Vertices are displayed where at least two subjects met the reliable-voxel criterion. Color indicates the mean held-out Spearman rank correlation between voxel response and the real-world size of the primary object, averaged across contributing subjects. Blue indicates greater responses to large objects, and red indicates greater responses to small objects. See Figure 3 for individual-subject maps and ROI overlays. **(B)** Proportion of voxels in each ROI classified as reliably small-preferring, reliably large-preferring, or neither. **(C)** Mean held-out Spearman rank correlation across all voxels in each ROI. Negative values indicate greater responses to small objects, and positive values indicate greater responses to large objects. Asterisks indicate significant across-subject effects after Benjamini–Hochberg FDR correction across ROIs (*q <* 0.05). In **(B)** and **(C)**, bars and error bars show mean *±*1 SEM across subjects, and circles indicate individual subjects.

**Figure 3:**
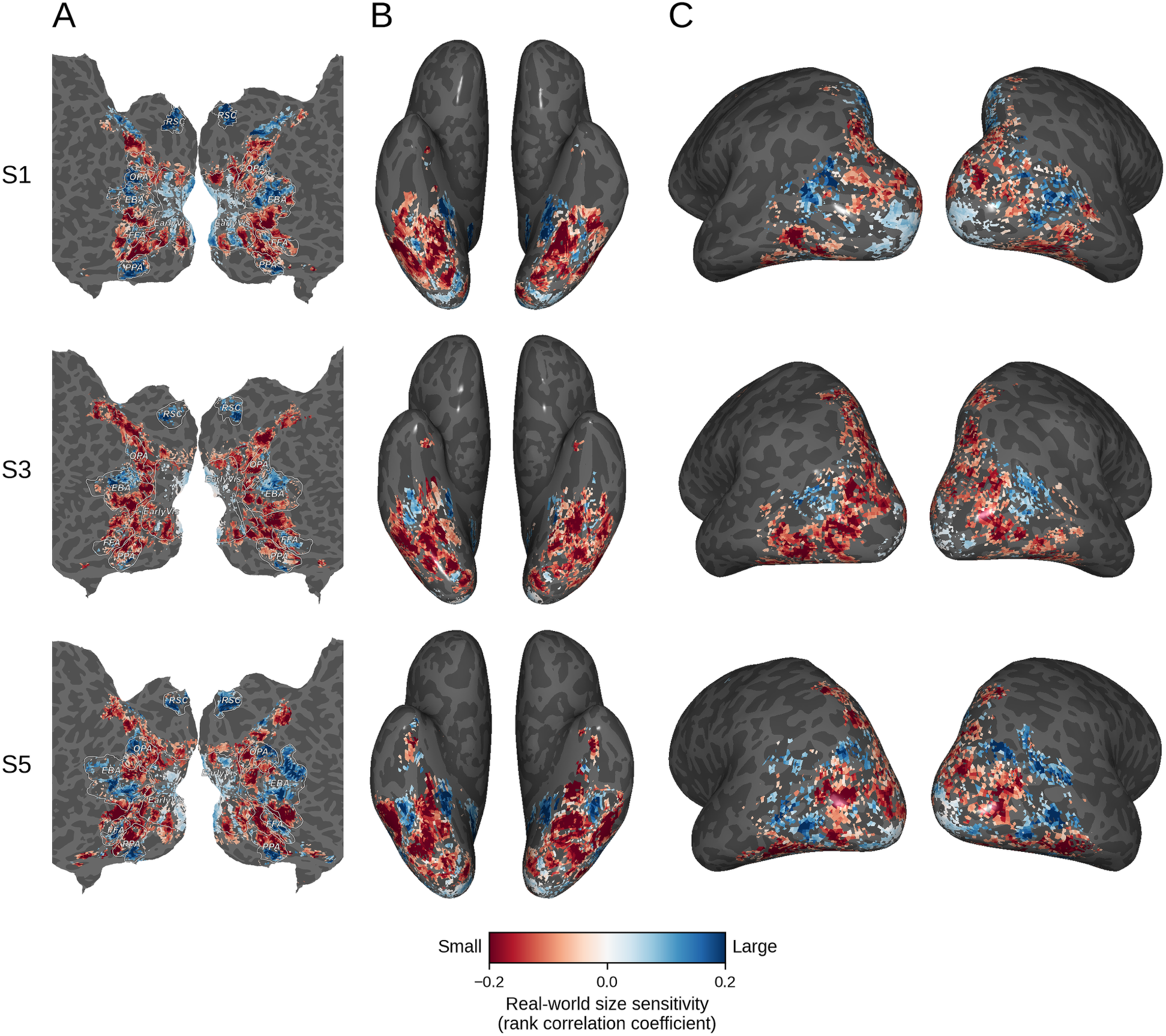
Cortical sensitivity to real-world object size in individual subjects. Size-sensitivity maps are shown for three example NSD subjects using the same reliable-voxel criterion and the same color scale as Figure 2A. Color indicates the mean held-out Spearman rank correlation, with blue indicating greater responses to large objects and red indicating greater responses to small objects. **(A)** Flatmap view with outlines of selected reference ROIs. **(B)** Ventral view. **(C)** Inflated lateral view. The three subjects show similar spatial organization, with variation in the precise location and extent of the effects.

Across all subjects, we found size-sensitive voxels on the ventral, lateral, and dorsal surfaces of occipitotemporal cortex. Along the ventral surface, this size sensitivity manifested as a spatially alternating pattern of small- and large-preferring voxels. Large-preferring voxels (blue) were prominent medially near the parahippocampal gyrus (partly overlapping areas PPA and RSC), as well as in a small patch along the fusiform gyrus (partly overlapping FFA), while small-preferring voxels (red) were present across adjacent ventral temporal cortex, including the region between the fusiform and parahippocampal gyri and more lateral territory near the inferior temporal gyrus. The individual-subject maps show a broadly similar alternating spatial pattern across the ventral surface, with variation in the precise location of the small- and large-selective regions across subjects.

Along the lateral surface of occipital cortex, we also observed alternating patches of small- and large-preferring voxels. Large-preferring responses were prominent in portions of lateral occipitotemporal cortex, including regions overlapping EBA, while small-preferring responses were seen across a wide region of dorsal cortex extending into the intraparietal sulcus. Early retinotopic visual regions (V1–hV4) contained intermingled small- and large-preferring voxels, with slightly more small-preferring voxels overall and some large-preferring responses present in more posterior (foveal) portions of early visual cortex.

We used functionally defined ROIs to further quantify these results, which showed that the proportions of small- and large-preferring voxels differed markedly across ROIs (Fig. 2B). RSC contained the largest proportion of large-preferring voxels among the examined ROIs. Within RSC, large-preferring voxels substantially outnumbered small-preferring voxels. In contrast, PPA and OPA, the other scene-selective ROIs, contained more small-than large-preferring voxels.

Word-selective region VWFA contained the largest proportion of small-preferring voxels among the examined ROIs, with substantially more small-than large-preferring voxels. OWFA also contained more small-than large-preferring voxels, although the majority of OWFA voxels lacked significant size sensitivity. Similar to the word-selective regions, face- and body-selective regions (EBA, FBA, OFA, FFA) all contained more small-preferring than large-preferring voxels, with FBA having the second-largest proportion of small-preferring voxels among all ROIs.

Computing the average correlation coefficients across voxels within each of these ROIs (Fig. 2C; Table 1), we found significantly greater responses to large objects in RSC and significantly greater responses to small objects in V1–hV4, FBA, and VWFA (see Table 1 for test statistics). None of the remaining ROIs showed significant mean size sensitivity after FDR correction (*q <* 0.05).

**Table 1:** ROI-averaged real-world size sensitivity (Fig. 2C). One-sample *t*_7_ statistics across the eight subjects and Benjamini–Hochberg FDR-corrected *q* values are reported across the ten ROIs. Positive *t* values indicate greater responses to large objects, whereas negative *t* values indicate greater responses to small objects.

| ROI | $t_7$ | $q$ | ROI | $t_7$ | $q$ |
| --- | --- | --- | --- | --- | --- |
| V1–V4 | −8.547 | 0.0002 | FBA | −6.747 | 0.0007 |
| RSC | 8.498 | 0.0002 | OFA | −2.255 | 0.0979 |
| PPA | 0.337 | 0.7460 | FFA | −2.273 | 0.0979 |
| OPA | −1.888 | 0.1262 | OWFA | −1.926 | 0.1262 |
| EBA | −0.381 | 0.7460 | VWFA | −9.544 | 0.0002 |

To assess the generality of these results, we next examined size sensitivity using the same analysis method applied to the MS COCO subset of BOLD5000, an independently acquired natural image fMRI dataset (Fig. 4; see *Methods* for details).

**Figure 4:**
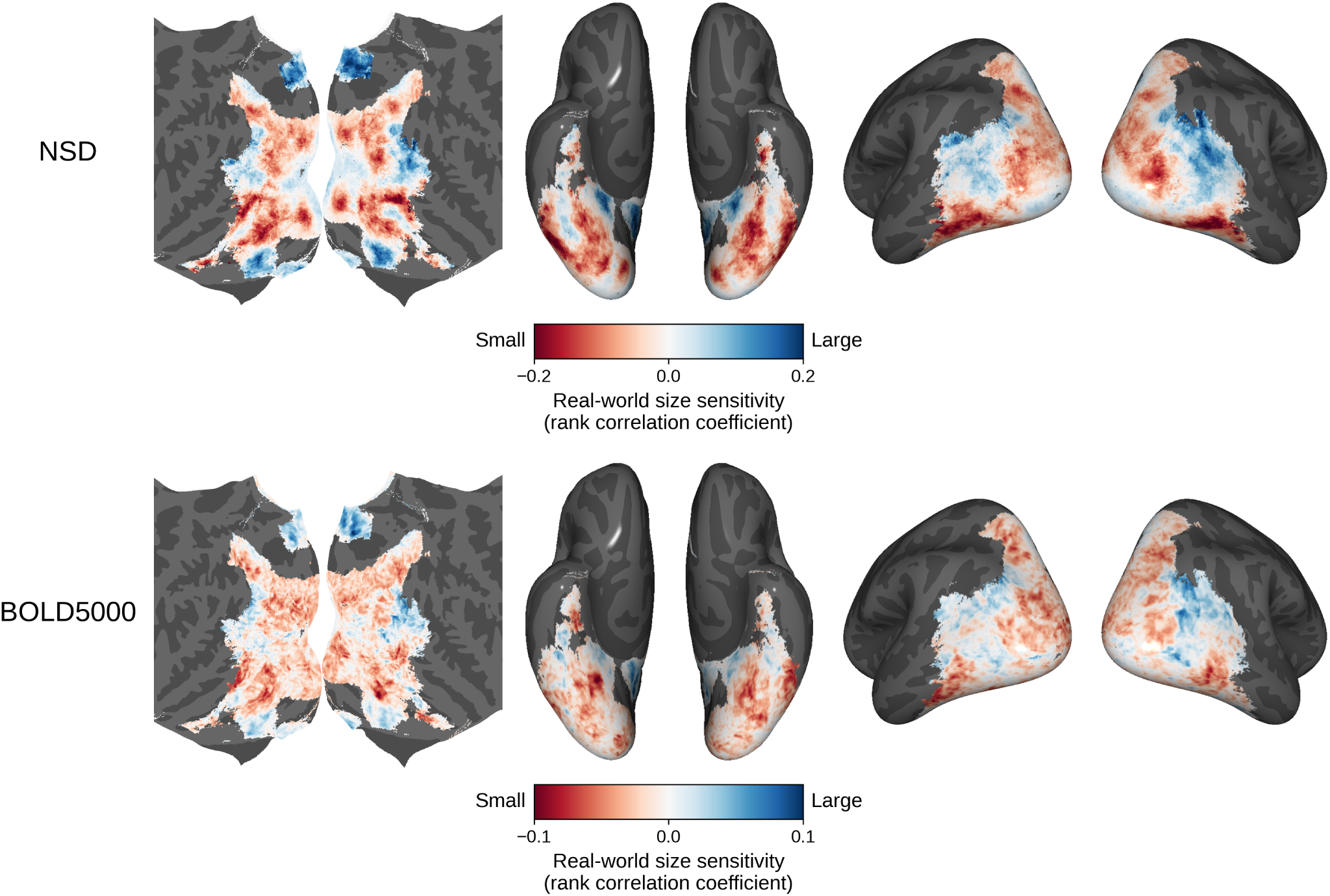
Cross-dataset comparison of real-world size sensitivity in NSD and BOLD5000. Group-average size-sensitivity maps from NSD and the MS COCO subset of BOLD5000 were projected to a common fsaverage surface. Size sensitivity was quantified at each vertex as the held-out Spearman rank correlation between fMRI response and the category-level real-world size estimate assigned to the primary object. From left to right, maps are shown in flatmap, ventral, and lateral views. Blue indicates greater responses to large objects, and red indicates greater responses to small objects. Both datasets are displayed on a common cortical support comprising vertices with data from at least two subjects in each dataset. For visualization purposes, NSD and BOLD5000 data are plotted on different color scales, with correlation ranges of *±*0.2 and *±*0.1, respectively.

We projected the NSD and BOLD5000 group-average size-sensitivity maps to a common fsaverage surface and compared their spatial organization across a common cortical support comprising vertices with data from at least two subjects in both datasets. At the group level, the BOLD5000 map showed spatial organization broadly similar to the NSD map, although the magnitude of size sensitivity values was smaller when computed using BOLD5000 versus NSD. Across the common cortical support, the NSD and BOLD5000 maps were positively correlated (spatial Pearson *r* = 0.64), with 69% sign agreement across vertices (*n* = 64,057). This correspondence was similar when the BOLD5000 analysis was restricted to COCO images shared with NSD (spatial Pearson *r* = 0.60; sign agreement = 67%) or to COCO images unique to BOLD5000 (*r* = 0.59; sign agreement = 67%; Supplementary Fig. S1), indicating that the cross-dataset correspondence was not restricted to shared stimuli. Individual BOLD5000 subject maps are shown in Supplementary Fig. S2.

We additionally examined the ImageNet stimuli in BOLD5000 to assess whether a similar organization was evident beyond MS COCO (Supplementary Fig. S3). The COCO and ImageNet BOLD5000 group maps showed moderate spatial correspondence across their full common cortical support (spatial Pearson *r* = 0.49; sign agreement = 61.0%). The ImageNet analysis showed a broadly similar but sparser pattern, with particularly reduced size sensitivity in early visual cortex.

These analyses showed group-level spatial correspondence in real-world size sensitivity across independently acquired natural-image datasets, while also revealing differences in the strength of the effect across stimulus sets. Comparisons with previous work using isolated object images are considered in the Discussion.

### 3.2 Analysis 2: Size sensitivity for animate and inanimate objects

Analysis 1 establishes the presence of real-world object size sensitivity across visual cortex.

The ROI results further show that the direction of size sensitivity differs across category-selective regions. Next, we assessed whether the spatial organization of size sensitivity was also preserved across a functional distinction that is known to elicit differential responses across these regions: animate versus inanimate objects. This question was partially motivated by past work (Konkle & Caramazza, 2013) in which size sensitivity was found for inanimate objects, but not for animate objects.

To test this, we separately estimated held-out size sensitivity for animate and inanimate object images using the set of voxels identified as reliably size-sensitive in Analysis 1 (Fig. 5; see *Methods* for details). We found voxel-wise size sensitivity to be broadly similar across animate and inanimate images, with both subsets showing an alternating spatial pattern of small- and large-preferring responses (Fig. 5C). Both maps broadly resembled the spatial organization observed in Analysis 1 (Figure 2A). A notable difference between the animate and inanimate maps, however, was that the bilateral large-preferring responses near the fusiform region (which were observed in Analysis 1) were substantially less pronounced in the animate map than in the inanimate map.

**Figure 5:**
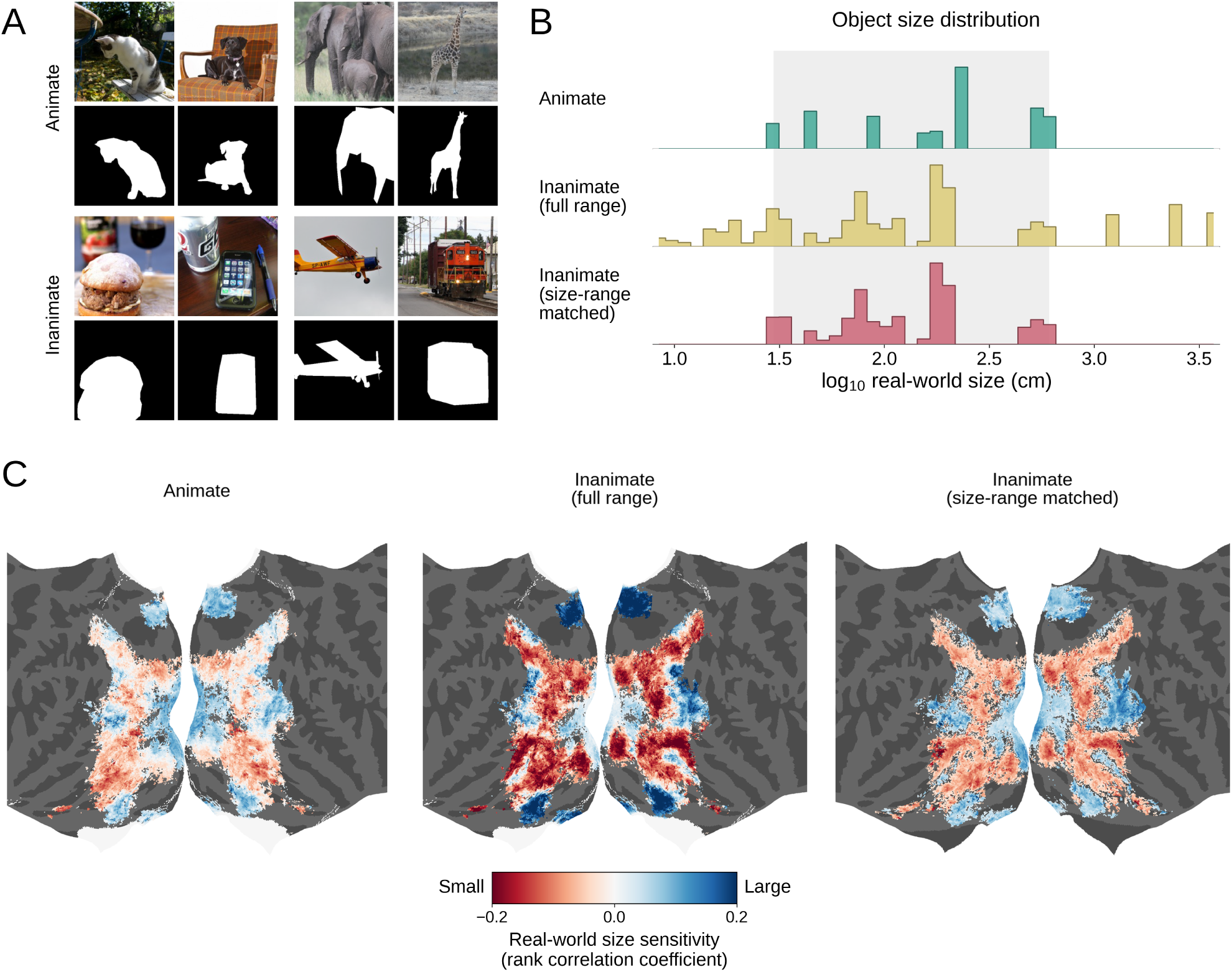
Comparison of size sensitivity for animate and inanimate objects. **(A)** Examples of animate (top) and inanimate (bottom) stimuli, shown with the mask of the most salient object. **(B)** Distributions of the image-level real-world size labels for animate images, all inanimate images, and inanimate images restricted to the animate size range. Sizes are shown in log_10_ centimeters. The light gray band marks the central animate size range (2.5th–97.5th percentiles) used to define the size-range-matched inanimate subset. **(C)** Group size-sensitivity maps for animate images, all inanimate images, and size-range-matched inanimate images. Voxel reliability was defined using all valid images, rather than separately within each subset, using the same reliable-voxel criterion as in Figure 2. Color indicates the mean held-out Spearman rank correlation across folds, averaged across subjects meeting the reliability criterion at each vertex. Maps are displayed on a common cortical support comprising vertices with at least two subjects meeting the reliability criterion. Blue indicates greater responses to large objects, and red indicates greater responses to small objects.

On average, a proportion of 0.7899 (SEM= 0.0248) of the reliably size-sensitive voxels across subjects had the same direction of size sensitivity (i.e., the same sign of the correlation coefficient) for animate and inanimate images. The average spatial Pearson correlation between the animate and inanimate size-sensitivity maps was *r* = 0.5768 (SEM= 0.0248).

Despite this spatial correspondence, the overall magnitude of size sensitivity was larger for the inanimate stimulus set. Across voxels, the mean absolute value of the held-out Spearman correlation was smaller for animate images (M = 0.0641, SEM= 0.0022) than for inanimate images (M = 0.1237, SEM= 0.0041), consistent with the visibly greater magnitude of both small- and large-preferring responses in the full-range inanimate map (Fig. 5C).

We first asked whether this difference could simply reflect the smaller number of animate images. Randomly downsampling the inanimate images to match the number of animate images left the magnitude of inanimate size sensitivity essentially unchanged (mean absolute held-out Spearman correlation: *M* = 0.1237, SEM= 0.0041). Animate–inanimate map correspondence was also nearly unchanged after count matching (mean within-subject spatial Pearson *r* = 0.5764, SEM= 0.0248; sign agreement = 0.7891, SEM= 0.0249). Thus, the weaker magnitude of size sensitivity for animate images was not attributable to the smaller number of animate stimuli.

By contrast, as shown in Fig. 5B, the animate images spanned a substantially narrower range of real-world sizes. Restricting the inanimate images to match the size range of animate images (Fig. 5C, right) reduced the mean magnitude of size sensitivity (absolute Spearman correlation) to 0.0735 (SEM= 0.0037), substantially closer to the magnitude observed for animate images, with both small- and large-preferring responses attenuated relative to the full-range inanimate map. Importantly, the size-range-matched inanimate map preserved the broad alternating spatial organization observed with the full range of inanimate images.

These analyses indicated that animate and inanimate images produced broadly similar spatial organization of real-world size sensitivity, despite larger correlation magnitude in the full-range inanimate stimulus set. The substantial attenuation of this magnitude difference after matching the represented size range suggested that much of the difference between size sensitivity for animate and inanimate objects was associated with the broader range of real-world sizes encompassed by the inanimate images.

### 3.3 Analysis 3: Robustness to feature and object-level controls

A recurring challenge in interpreting cortical selectivity for high-level visual properties such as real-world size is the extent to which they are driven by low or mid-level visual features as opposed to high-level semantic representations (Andrews et al., 2015; Julian et al., 2017; Long et al., 2018; Nasr et al., 2014). Because small and large objects may differ systematically in their image statistics (e.g., spatial frequency, the prevalence of rectilinear versus curved contours), we asked how much of the observed size sensitivity remained after accounting for such visual features. Namely, we constructed voxelwise linear models based on Gabor, GIST, and sketch token features in combination with an estimate of each voxel’s population receptive field (pRF). As described in the *Methods*, these three feature spaces represent local low-level, global low-level, and mid-level features, respectively (Fig. 6). We first evaluated the held-out variance explained (*R*^2^) by each feature model (Fig. 6B). All three models explained the greatest variance in early visual cortex. Sketch token features explained relatively more variance than Gabor features in some higher visual regions, whereas GIST explained relatively more variance in RSC.

**Figure 6:**
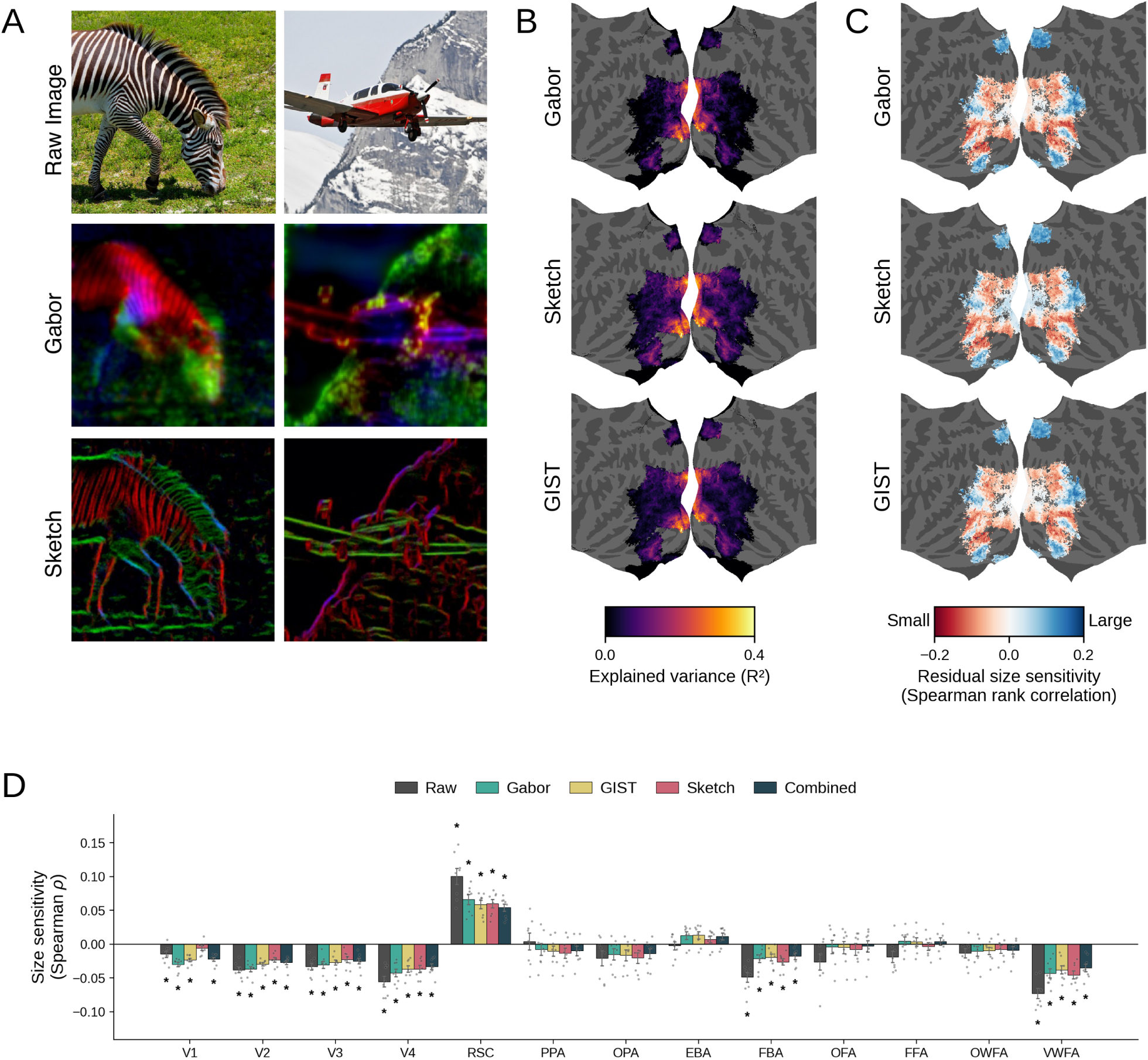
Controlling for low- and mid-level visual features. **(A)** Example NSD images with corresponding Gabor and sketch-token feature representations. Feature representations depict PCA projections of the feature maps from each model; see *Methods* for details on all feature models. **(B)** Group-average held-out explained variance (*R*^2^) for the Gabor, sketch-token, and GIST models, shown on a flattened fsaverage surface. **(C)** Group size-sensitivity maps after controlling for each feature space. Color indicates the residual held-out Spearman rank correlation. Maps are displayed on the same reliability-defined cortical support used for the primary size-sensitivity map (Fig. 2). Blue indicates greater responses to large objects, and red indicates greater responses to small objects. **(D)** ROI-level size sensitivity before feature control and after controlling for Gabor, GIST, sketch-token, or all three feature spaces combined. Bars and error bars indicate mean *±*1 SEM across subjects, and gray points indicate individual subjects. Asterisks indicate significant residual size sensitivity after Benjamini–Hochberg FDR correction (*q <* 0.05) across the ROI-by-feature-model tests.

We next recomputed held-out size sensitivity after accounting for each feature space using the feature-control procedure described in the *Methods* (Fig. 6C). The large-scale distribution of small- and large-object preferences remained visible after controlling for each feature space, although the magnitude of size sensitivity was generally reduced. At the ROI level, the large-object preference in RSC and the small-object preferences in FBA and VWFA were each smaller in magnitude but retained significance after controlling for every low- and mid-level feature space (Fig. 6D). Our previous analysis (Fig. 2) had also indicated a significant preference for small objects in V1–hV4, so to better understand this effect, here we individually analyzed results in V1, V2, V3, and hV4. This revealed that the magnitude of preference for small objects was smallest in V1 and increased progressively from V2 to V3 to hV4. In V2–hV4, this preference for small objects was attenuated by each of the feature controls but retained significance. In V1, size sensitivity retained significance following the Gabor and GIST controls but was not significant following the sketch token control. This suggests that mid-level features may account for some of the size sensitivity observed in V1.

Although these low- and mid-level feature controls reduced the magnitude of size sensitivity in several ROIs, its spatial organization was largely preserved. Across subjects, the map obtained after jointly controlling for all three feature spaces remained strongly aligned with the original size-sensitivity map (mean within-subject spatial Pearson *r* = 0.941, SEM= 0.004; sign agreement = 0.915, SEM= 0.012). Similar spatial correspondence was observed for the individual feature controls (Supplementary Table S1). Thus, the tested low- and mid-level visual features attenuated size sensitivity but did not fully account for its cortical distribution.

As a complementary encoding analysis, we tested whether real-world size provided predictive information about voxel responses beyond the combined visual feature representation. Adding real-world size to the combined feature model produced a positive mean increase in held-out explained variance in every ROI. This increase was significant after FDR correction (*q <* 0.05) in 12 of the 13 ROIs, with OWFA as the only ROI that did not reach significance (Supplementary Table S2). Although the increases were small in absolute magnitude, they indicated that real-world size provided incremental predictive information beyond the tested visual feature spaces.

We next tested whether the results depended on how the image-level size label was defined when an image contained multiple objects (Fig. 7A–B). We recomputed size sensitivity using the saliency-primary object, the largest object, an area-weighted summary, and a saliency-weighted summary. V2–hV4, RSC, FBA, and VWFA remained significantly size-sensitive under all four definitions. V1 remained significant under the saliency-primary, largest-object, and saliency-weighted definitions, but not under the area-weighted definition. Both weighted definitions additionally yielded significant small-object preferences in EBA, OFA, FFA, and OWFA. No significant ROI-level effect reversed direction across the object-definition analyses.

**Figure 7:**
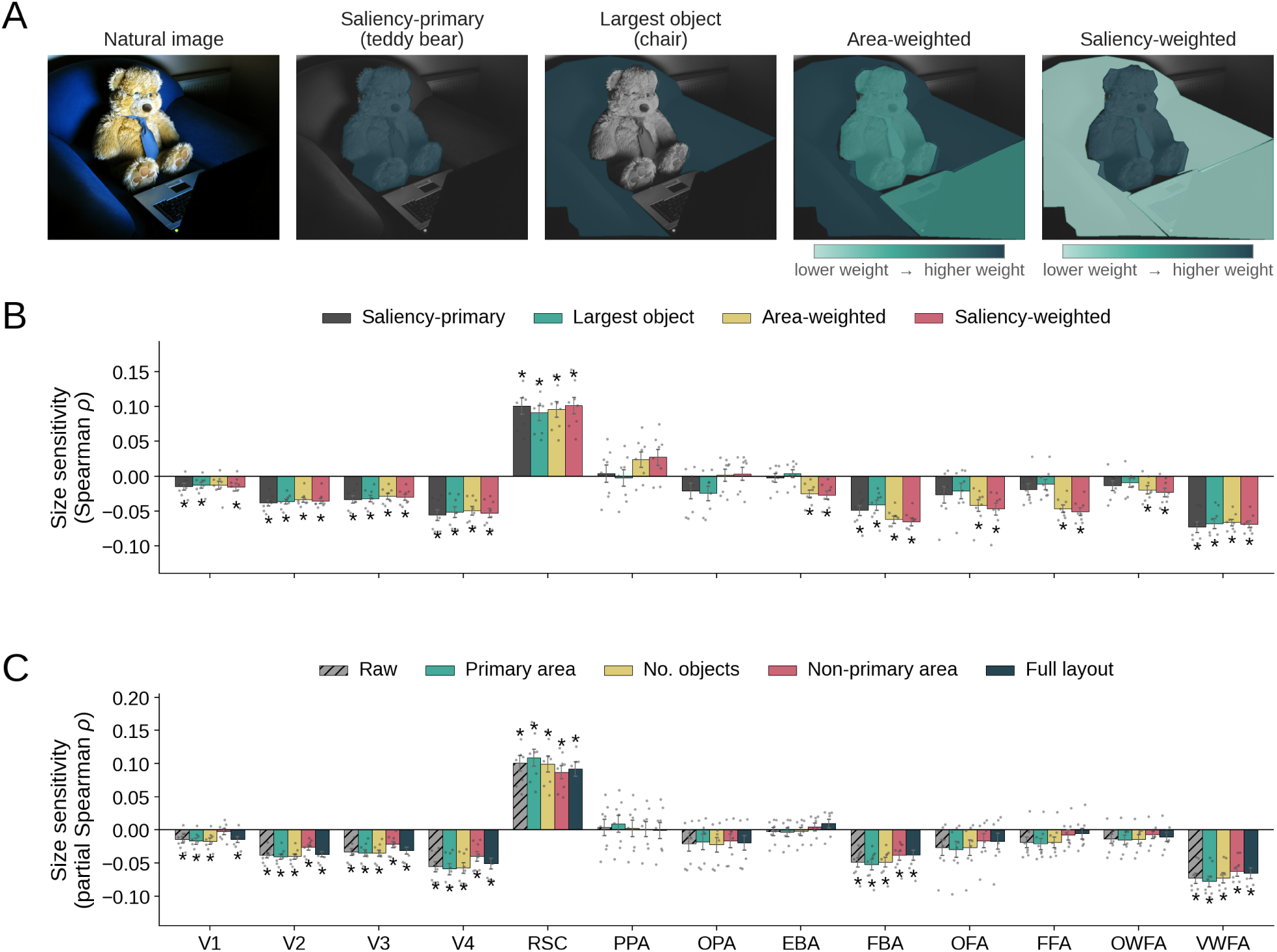
Robustness of real-world size sensitivity to object-definition and object-layout controls. **(A)** Illustration of four rules for defining the image-level real-world size label: saliency-primary, largest-object, area-weighted, and saliency-weighted. For the weighted definitions, mask intensity indicates each object’s normalized contribution to the resulting size label. **(B)** ROI-level size sensitivity estimated using each object-definition rule. **(C)** ROI-level size sensitivity before object-layout control (raw) and after controlling for primary-object area, number of objects, non-primary-object area, or all layout variables jointly (Full layout). Size sensitivity in Panels B and C was quantified using Spearman rank correlation. Bars and error bars indicate mean *±*1 SEM across subjects, and gray points indicate individual subjects. Asterisks indicate significant size sensitivity after Benjamini–Hochberg FDR correction (*q <* 0.05) across the ROI-by-control tests within each control family. The raw condition in Panel C is shown for reference and was not included in the object-layout correction family.

Finally, we tested whether size sensitivity remained after accounting for coarse properties of object layout and scene clutter (Fig. 7C). We separately controlled for the area of the primary object, the number of objects, the total image area occupied by non-primary objects, and all three layout variables jointly. After FDR correction (*q <* 0.05), V2–hV4, RSC, FBA, and VWFA remained significantly size-sensitive under every layout control. V1 remained significant after controlling for primary-object area, object count, and all three layout variables jointly, but not after controlling for non-primary-object area. No ROI changed its direction of preference under any layout control. Across all four layout controls, the controlled maps also remained strongly aligned with the original size-sensitivity map. For example, the joint layout control yielded a mean within-subject spatial Pearson correlation of *r* = 0.9873 (SEM= 0.0005) and a mean sign agreement of 0.9925 (SEM= 0.0011) across subjects (see Supplementary Table S1 for all test values).

These analyses showed that the principal ROI-level patterns of real-world size sensitivity were generally preserved across the tested visual-feature, object-definition, and object-layout analyses, although the magnitude and statistical strength of individual effects varied across controls. Moreover, the pattern of size sensitivity across all individual voxels was also largely preserved across these controls, underscoring the reliability of these topographic size-sensitivity maps.

### 3.4 Analysis 4: Real-world size tuning profiles

Prior work has explored size sensitivity using objects discretely categorized as large or small. However, real-world object sizes span several orders of magnitude. This raises the question of whether the relationship between object size and neural response (i.e., the shape of size-tuning profiles) differs across large-versus small-preferring populations. To characterize the shape of this relationship, we compared a set of candidate monotonic functions relating normalized real-world size to voxel response (Fig. 8A).

**Figure 8:**
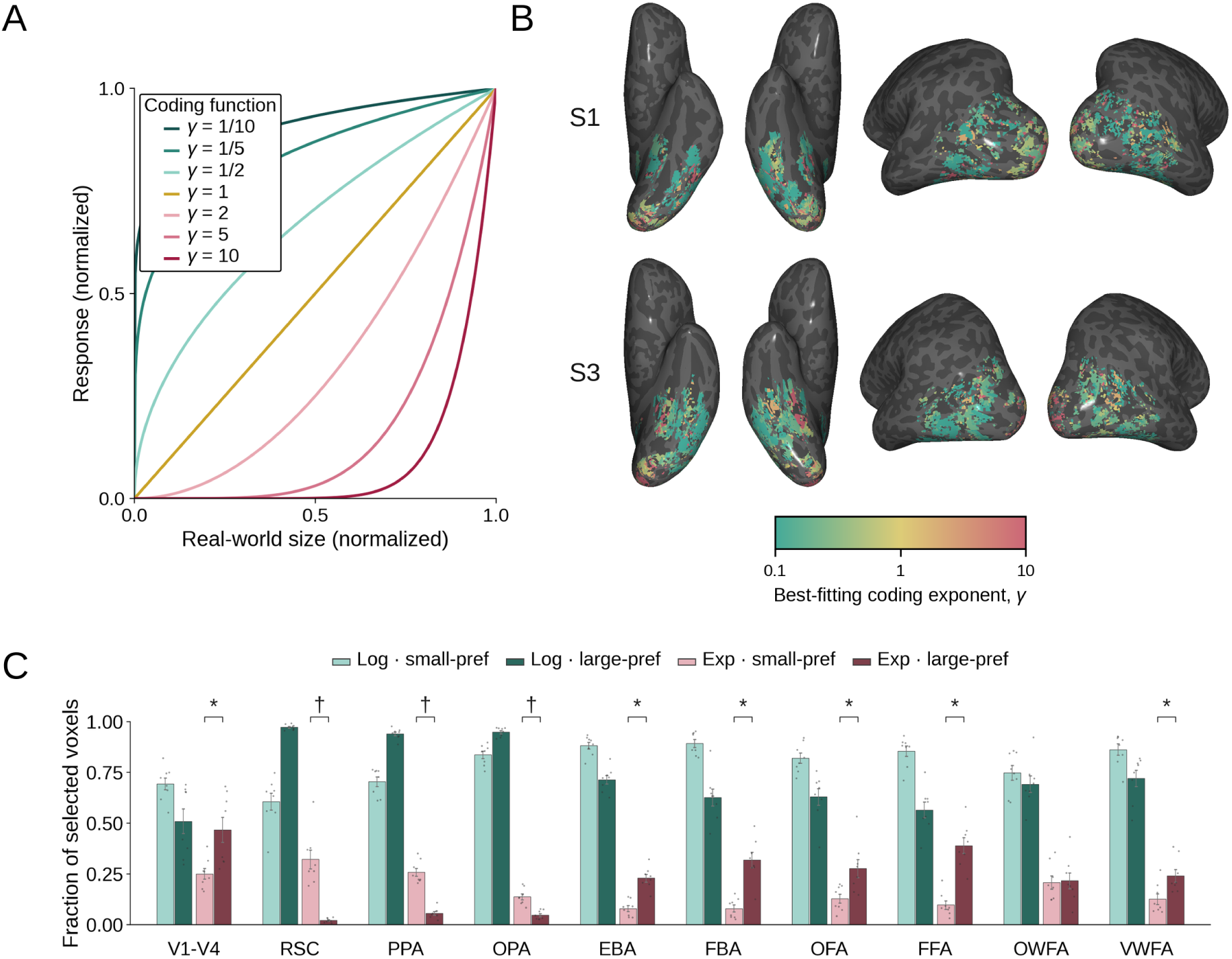
Real-world size tuning profiles. **(A)** Candidate power functions of the form *f* (*x*) = *x^γ^*, relating normalized real-world size to neural response. Values of *γ <* 1, *γ* = 1, and *γ >* 1 correspond to logarithmic-like, linear, and exponential-like functions, respectively. **(B)** Best-fitting coding exponent *γ* for size-sensitive voxels in two example subjects. Voxels are shown within the ROIs included in the tuning analysis if they met the reliable size-sensitivity criterion; color indicates the median best-fitting exponent across the folds in which each voxel was selected. **(C)** Fractions of size-sensitive small- and large-preferring voxels for which logarithmic-like or exponential-like functions provided the best fit among the tested candidates, shown by ROI, with the two preference groups displayed together within each ROI. Linear fits are omitted for clarity. Bars and error bars indicate mean *±*1 SEM across subjects, and gray points indicate individual subjects. Asterisks indicate a greater exponential-like fraction among large-than small-preferring voxels, whereas daggers indicate the opposite pattern (paired two-sided *t*-tests with Benjamini–Hochberg FDR correction (*q <* 0.05) across ROIs).

For functions taking the form of *f* (*x*) = *x^γ^*, we searched through logarithmic-like (*γ <* 1), linear (*γ* = 1), and exponential-like (*γ >* 1) functions (see *Methods* for details).

Logarithmic-like functions expand differences among smaller sizes while compressing differences among larger sizes, while exponential-like functions show the opposite pattern. We therefore asked whether the best-fitting transformation differed between small- and large-preferring voxels.

Using the cross-validated procedure described in the *Methods* to identify reliably size-sensitive voxels, we identified within each fold the candidate value of *γ* that yielded the highest held-out *R*^2^ for each selected voxel (Fig. 8B).

Across selected voxel fits, logarithmic-like functions were most frequently selected overall (76.1%), followed by exponential-like (19.7%) and linear (4.3%) functions.

Within each ROI, we separately tested whether logarithmic-like functions accounted for more than half of the selected small-preferring and large-preferring voxels. Logarithmic-like functions constituted a significant majority in 18 of these 20 comparisons after FDR correction (*q <* 0.05); the only exceptions were large-preferring voxels in pooled V1–hV4 and FFA (Fig. 8C).

We next tested whether exponential-like fits were differentially represented among small- and large-preferring voxels within each ROI. Exponential-like fits were significantly more frequent among large-preferring voxels versus among small-preferring voxels in pooled V1–hV4, EBA, FBA, OFA, FFA, and VWFA after FDR correction (*q <* 0.05). This enrichment was particularly pronounced in FFA, where the mean exponential-like fraction increased from 0.097 among small-preferring voxels to 0.390 among large-preferring voxels. In contrast, RSC, PPA, and OPA showed the opposite pattern, with exponential-like fits more frequent among small-preferring than large-preferring voxels, whereas OWFA showed no significant difference. Thus, although logarithmic-like functions were selected most frequently overall, exponential-like fits showed region-dependent enrichment across small- and large-preferring voxel populations.

## 4 Discussion

Real-world object size is a behaviorally relevant property that guides interactions with both objects and the environment. Given that estimates of real-world size often depend on context, we explored how using complex natural scene images as stimuli affects cortical responses. Across multiple analyses, reliable topographic patterns for real-world size sensitivity were observed in higher visual cortex. Moreover, these patterns were observed with both animate and inanimate objects, and consistent size preferences were observed for several category-selective ROIs. These findings provide new insights into how the physical size of objects is cortically represented during naturalistic vision.

In our study, size sensitivity manifested as alternating regions of sensitivity for small and large objects along the ventral and lateral surface of higher visual cortex. This organization is aligned with the patterns observed in earlier fMRI experiments studying the neural encoding of real-world size using isolated objects (Julian et al., 2017; Konkle & Caramazza, 2013; Konkle & Oliva, 2012). However, one difference from Konkle and Caramazza (2013) is our observation of bilateral large-preferring regions adjacent to the fusiform region (Figure 3) along with small-preferring regions medial to these; importantly, this pattern was observed using both NSD and BOLD5000 and was particularly evident for inanimate images (Figure 5C). A similar pattern in ventral cortex has been reported for the THINGS dataset (Hebart et al., 2023; Stoinski et al., 2026).

The fact that the THINGS images are generally more object-centric and less scene-rich as compared to MS COCO images used in our study suggests that the patterns we observed are robust to differences in stimulus set properties. However, our results and those obtained using THINGS diverge in early retinotopic visual cortex, where we observe a small bias for large objects in center-preferring (posterior) voxels while analyses using THINGS show a bias for small objects in center-preferring voxels (Hebart et al., 2023; Stoinski et al., 2026); we speculate that this discrepancy may be due to stimulus set differences (e.g., the resultant degree of foveation by participants).

Reinforcing that stimulus set content might drive some of the neural responses to real-world size, our analysis of the ImageNet images in BOLD5000 revealed that COCO images give rise to an organization that corresponds closely to the results obtained using NSD, whereas the ImageNet images give rise to a similar but substantially sparser pattern, particularly in early visual cortex. Content differences across these image datasets include richness of scene context, object framing, category sampling, and the represented range of real-world sizes. Despite these differences, the overall cortical patterns for real-world size sensitivity converge across NSD, BOLD5000, and THINGS, suggesting that a size-based organization in visual cortex is not specific to a single image set.

This correspondence across datasets is notable given the importance of context in judgments of object size. For example, in the Ebbinghaus illusion, an object appears relatively larger when surrounded by small objects, but relatively smaller when surrounded by large objects (Doherty et al., 2010). Similar observations have been made using the Jastrow (Agrillo et al., 2019), Delboeuf (Surkys et al., 2006), and vertical-horizontal (Künnapas, 1955) illusions, in which perceived size is affected by contextual cues. One interpretation of this consistency across both isolated object and natural scene stimuli is that context becomes increasingly important as the real-world size of an object becomes more ambiguous or entirely unknown. However, due to prior knowledge, non-contextual cues to size, and other factors, in many natural images real-world size estimates are accessible without taking into account context. At the same time, differences in the contextual richness of stimulus images may contribute to the novel patterns observed both in the present work and in THINGS, underscoring the possibility that a more complete picture of human cortical function may emerge from incorporating both controlled and naturalistic stimuli.

At a macro level, ROI analyses help relate the observed voxelwise organization to functionally defined cortical systems. However, we note that ROI-level averages do not imply that all voxels within a region share the same preference. As shown in the flatmaps (Fig. 3A), individual ROIs can contain both small- and large-preferring regions, whereas the ROI-level statistic only summarizes the net effect across the entire voxel population. Pooled V1–hV4 provides one example: early visual cortex contains intermingled small- and large-preferring voxels, yet shows a significant small-preferring effect at the ROI-averaged level.

The same caveat should be taken into account when interpreting the scene-selective ROIs. PPA and OPA each contain local populations of both small- and large-preferring voxels, with slightly more reliably small-preferring voxels overall, but neither region shows a significant net preference when responses are averaged across all ROI voxels. RSC, by contrast, shows a significant large-preferring effect and contains the greatest proportion of reliably large-preferring voxels among examined ROIs. This heterogeneous pattern across scene-selective regions differs from Konkle and Caramazza (2013) and Julian et al. (2017), who reported large-object preferences across all three ROIs. Our conjecture is that differences in the size sensitivity of PPA and OPA may be due to the use of scene stimuli as opposed to isolated objects, or to the range of real-world sizes used across experiments.

Similarly, differences in size sensitivity within the same category-selective network appear in word- and body-selective cortex. VWFA and FBA show significant small-preferring effects, whereas OWFA and EBA do not reach significance at the ROI-averaged level. These contrasting preferences in RSC and VWFA may reflect the natural statistics of each visual category to which these regions are sensitive. In particular, scenes often contain large-scale environmental structures and support navigation (R. A. Epstein & Baker, 2019; Lescroart & Gallant, 2019), whereas words are typically encountered as relatively small-scale visual patterns (McCandliss et al., 2003). However, the differences between RSC and other scene-selective ROIs, between VWFA and OWFA, and between FBA and EBA indicate that this relationship is not uniform across regions within the same functional system. One possibility is that regions within the same system emphasize different aspects of the preferred category or differ in their sensitivity to spatial and contextual information, making real-world size more impactful in some regions than in others.

Across both animate and inanimate objects, we observed sensitivity to real-world size, accompanied by corresponding cortical patterns across the two domains. This result contrasts with those of Konkle and Caramazza (2013), who reported size-dependent responses only within the inanimate domain. Similarly, much of the remaining literature on cortical size sensitivity has focused primarily on inanimate objects (Huang et al., 2022; Julian et al., 2017; Konkle & Oliva, 2012). However, studies using the THINGS dataset also observe size sensitivity within the animate domain (Stoinski et al., 2026). We posit that these cross-study differences are due to methodological differences such as the use of natural images and continuous size labels as opposed to discrete categories, or to recent improvements in fMRI signal quality. At the same time, we did find lower size sensitivity for animate versus inanimate objects, although the spatial patterns remained similar. Matching the inanimate images to the real-world size range spanned by the animate images substantially reduced this difference, suggesting that differences in the sampled size distributions may partly explain weaker animate effects and their absence in prior work.

Residual differences might reflect category sampling, within-category size variability, or other visual and semantic properties that differ between animate and inanimate images. Future work using stimulus sets with balanced size distributions across animacy domains, together with exemplar-level rather than category-level size estimates, could help establish whether this residual difference is associated with animacy itself or with correlated properties of the sampled stimuli.

As a control, we also investigated the contributions of low- and mid-level visual features to real-world object size sensitivity. In particular, real-world size may correlate with image statistics – for example, images containing large and small objects may differ in mid-level statistics such as boxiness or curvature (Long et al., 2016, 2018), or low-level statistics such as orientation or spatial frequency (Henderson et al., 2023).

Interestingly, there is some attenuation of size sensitivity when controlling for Gabor, GIST, and sketch token visual features, indicating that low- and mid-level image statistics contribute to the observed effects. Most notably, in V1, size sensitivity was not significant following control for mid-level sketch token features. At the same time, the broader spatial organization and direction of size preferences across higher visual cortex were largely preserved across our visual feature controls, and complementary analyses further established that real-world size provides predictive power beyond the tested visual features in nearly all examined ROIs. This pattern of results is consistent with prior evidence that coarse texture and form information do produce the large-scale cortical organization associated with real-world size, but with a weaker response magnitude relative to that elicited by intact recognizable stimuli (Long et al., 2018). We also tested the robustness of size sensitivity to additional controls, including alternative definitions of the primary object and image-layout covariates. Both analyses revealed a consistent spatial organization after accounting for these variables.

Together, these results suggest that low- and mid-level image statistics and object-layout properties account for some of the measured size sensitivity, but do not fully account for the overall magnitude of the effect or its consistent cortical organization. As such, the remaining size sensitivity may reflect information associated with object identity, semantic meaning, high-level visual features, and context that can be used to infer real-world size.

Finally, our tuning analysis provides some insight into how cortical response profiles vary across real-world object sizes. In earlier work, Konkle and Oliva (2011) found that the canonical visual size at which familiar objects are drawn, imagined, or preferred for viewing increases approximately logarithmically with their assumed real-world size. This relationship compresses the many orders of magnitude separating real-world objects into a more limited range of preferred visual sizes. The predominance of logarithmic-like response profiles in our analyses may reflect a related form of compression in cortical responses, with greater differentiation among smaller sizes and progressively less differentiation toward the larger end of the range. Although logarithmic-like profiles predominated overall, exponential-like fits were also observed, with their distribution varying across cortical systems. In early visual cortex and several category-selective regions, exponential-like fits were more common among large-preferring voxels, suggesting that responses changed most strongly at the upper end of the size range. Scene-selective cortex, however, was the notable exception. Across RSC, PPA, and OPA, exponential-like fits were rare among large-preferring voxels. Instead, the large-preferring voxels in scene-selective regions showed predominantly logarithmic-like profiles, suggesting strong differentiation between smaller and larger objects followed by increasing compression across the upper end of the size range. This pattern may relate to the role of scene-selective cortex in representing environmental structure and spatial information relevant for navigation (R. A. Epstein & Baker, 2019). One possibility is that these regions distinguish smaller objects from larger, navigation-relevant landmarks, while showing less differentiation among objects at the upper end of the size range. Future work should address how these differences in size coding across large- and small-preferring neural populations are related to psychophysical effects, human perception, and navigation behavior.

In conclusion, our findings reveal robust higher visual cortex sensitivity to the size of real-world objects in natural scenes. Importantly, a consistent organization was observed across independent natural-image datasets, was similar for animate and inanimate objects, and remained evident across controls for visual features, primary-object definition, and image layout. Interestingly, response profiles across the broad range of real-world object sizes differed according to preference direction and cortical region. Taken together, our findings provide new insight into how the functional organization of higher visual cortex supports our ability to extract behaviorally relevant object information from the environment.

## Data and code availability

The data used in this study have been made openly available by the authors of Allen et al. (2022) and Chang et al. (2019); see these original publications for details. The analysis codes are publicly available at https://github.com/ahmadhlayhel/cortical-real-world-size.

## Author Contributions

A.H. – Methodology, Software, Formal Analysis, Investigation, Visualization; Writing—Review & Editing; A.F.L. – Conceptualization, Data Curation, Methodology, Software, Formal Analysis, Investigation, Writing— Original Draft, Writing—Review & Editing, Visualization, and Project Administration. L.W. – Conceptualization, Writing—Review & Editing, Project Administration, Supervision. M.J.T. – Conceptualization, Writing—Review & Editing, Project Administration, Supervision. M.M.H. – Conceptualization, Data Curation, Software, Writing—Review & Editing, Project Administration, Supervision.

## Supplementary Material

### Prompt used to obtain category-level real-world size estimates

The following system and user prompts were used to query GPT-4o. The placeholder {category} was replaced with the name of each MS COCO category.

#### System prompt

You are a careful reference assistant that estimates the typical real-world physical dimensions of everyday objects. You describe a single common, full-size instance as encountered in ordinary daily life. You DO NOT describe toys, miniatures, scale models, decorative versions, packaging, or unusual extreme specimens, UNLESS the category name itself implies one (e.g., a “teddy bear” is a toy). For living things, describe a typical healthy adult. Give numeric estimates in centimeters. Respond with a single JSON object that matches the requested schema and nothing else.

#### User prompt

Estimate the typical real-world physical dimensions of a {category}.

Describe one common, full-size, real instance of `‘{category}’` as encountered in everyday life (for an animal or person, a typical adult). Exclude toys, miniatures, scale models, decorative or novelty versions, and unusually extreme specimens unless the word `‘{category}’` itself implies them.

Report, in centimeters:

- `length cm`: the typical longest horizontal extent
- `width cm`: the typical second horizontal extent
- `height cm`: the typical vertical extent

**Figure S1:**
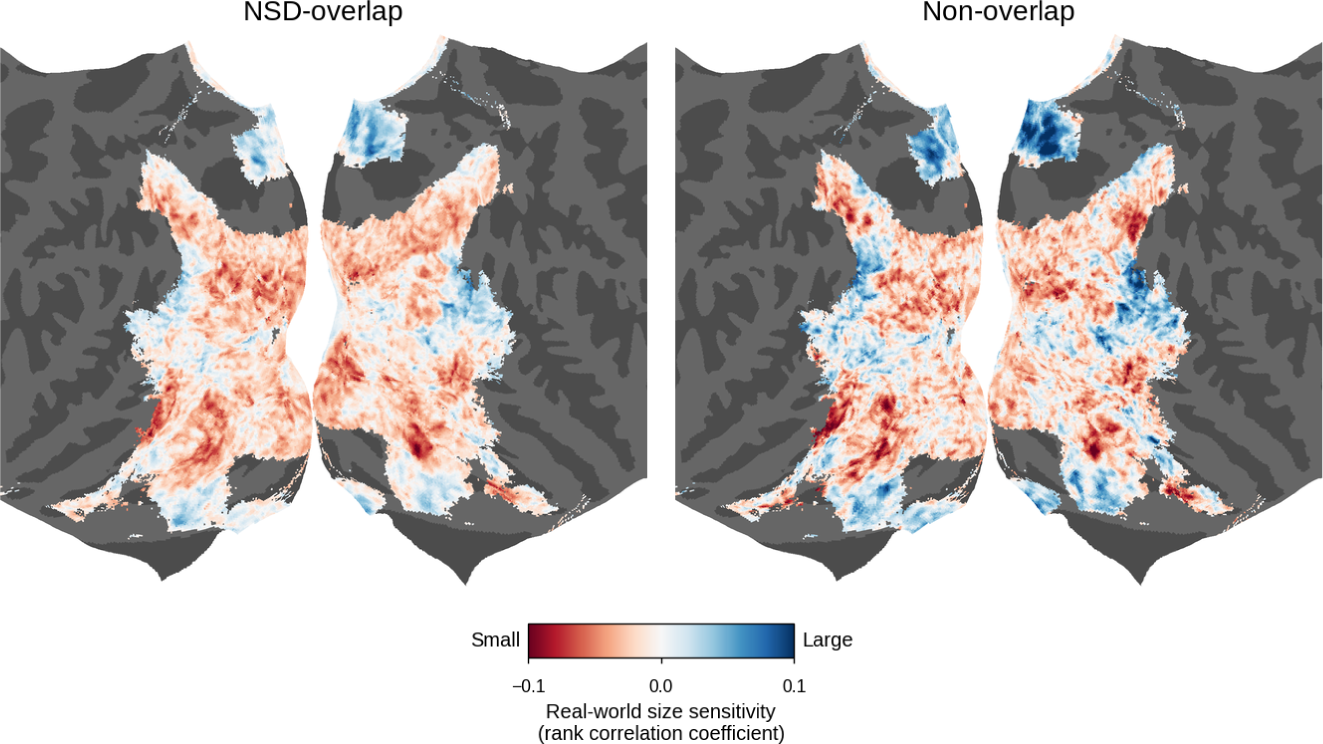
BOLD5000 size-sensitivity patterns for images overlapping with NSD and images unique to BOLD5000. BOLD5000 group maps were computed separately for COCO images shared with NSD and for COCO images unique to BOLD5000. Both maps are shown on the common cortical mask used in Figure 4 (*n* = 64,057 vertices). Agreement with the NSD group map was similar for the overlapping subset (spatial Pearson *r* = 0.60; sign agreement = 67%) and the non-overlapping subset (*r* = 0.59; sign agreement = 67%). Thus, the observed cross-dataset correspondence was not restricted to images shared between NSD and BOLD5000.

**Figure S2:**
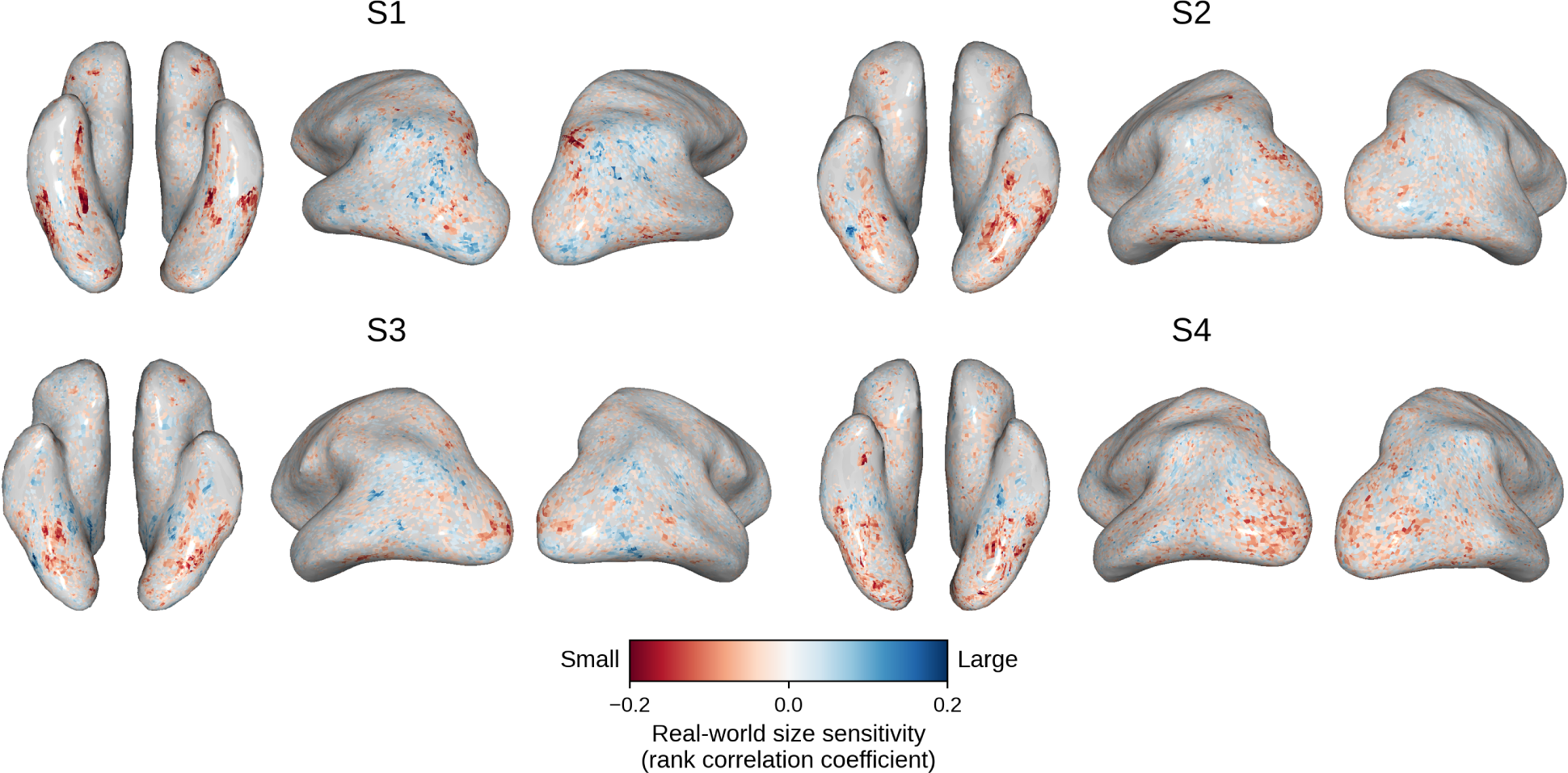
Individual-subject real-world size-sensitivity maps in BOLD5000. Size-sensitivity maps are shown for the four BOLD5000 subjects on their native cortical surfaces. Color indicates the Spearman rank correlation between voxel response and the real-world size label of the primary object, with blue indicating greater responses to large objects and red indicating greater responses to small objects. These maps show the individual-subject patterns underlying the BOLD5000 group comparison in Figure 4.

##### BOLD5000 ImageNet analysis

For the supplementary ImageNet analysis, each stimulus was assigned the real-world size of its original ImageNet target category, identified from the corresponding synset, rather than using the saliency-guided object-selection procedure applied to MS COCO images (because ImageNet does not include instance segmentation masks). The validity of this approach is supported by the robustness of the primary COCO results to alternative object-definition rules, which largely preserved the spatial organization and direction of size preferences. Real-world size estimates for the ImageNet categories were obtained using the same GPT-4o estimation procedure used for the COCO categories (see *Methods*), with ten estimates collected per category and the largest reported physical dimension averaged across responses before log transformation. Categories matching previously validated COCO labels reused those estimates. For the maps shown here, size sensitivity was estimated from held-out data using the same ten split-half partitions and both fold directions described in the primary analysis. SUN stimuli were not analyzed because they are labeled by scene category rather than object category and therefore do not provide a corresponding object-level label from which real-world size can be assigned.

**Figure S3:**
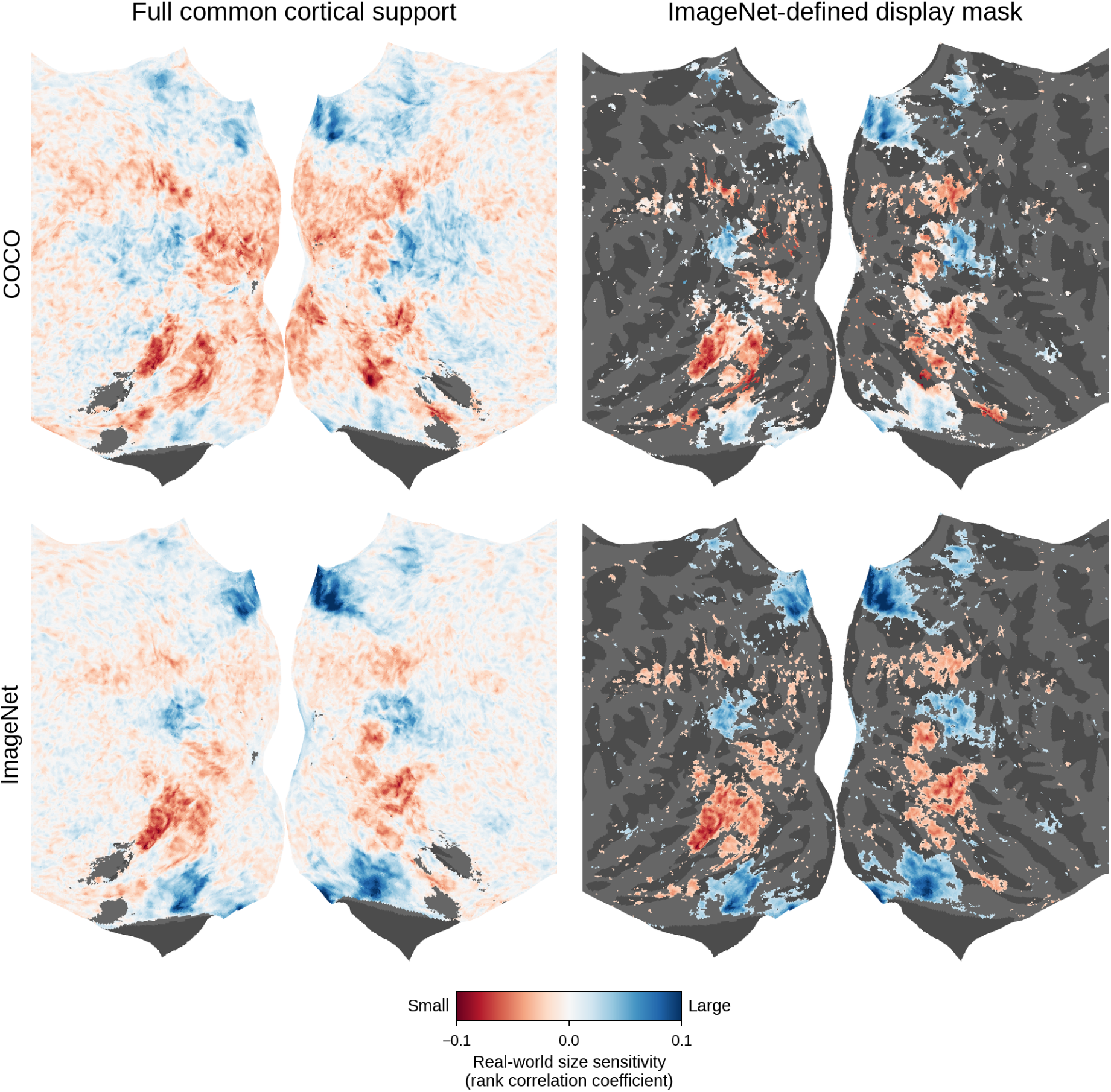
Comparison of real-world size sensitivity for COCO and ImageNet stimuli in BOLD5000. Group-average size-sensitivity maps were computed separately using the COCO (top row) and ImageNet (bottom row) stimulus subsets of BOLD5000. The left column shows the full common cortical support, comprising vertices with finite estimates in both maps (*n* = 295,946), with no correlation-magnitude threshold applied. Across this support, the COCO and ImageNet maps showed moderate spatial correspondence (spatial Pearson *r* = 0.49; sign agreement = 61.0%). The right column shows the same maps restricted to an ImageNet-defined display mask (*|ρ*_ImageNet_*| >* 0.02; *n* = 36,823 vertices). Within this mask, spatial correspondence was stronger (spatial Pearson *r* = 0.78; sign agreement = 84.9%). Blue indicates greater responses to large objects, and red indicates greater responses to small objects.

**Table S1:** Spatial correspondence after visual-feature and object-layout controls. Values report the across-subject mean *±* SEM of within-subject comparisons between the original and controlled size-sensitivity maps. All controls were evaluated over the same subject-specific set of reliably size-sensitive voxels, selected in at least 11 of the 20 cross-validation folds, within the shared set of focus ROIs. The mean number of evaluated voxels per subject was 6,698 (range: 4,903–8,196).

| Analysis family | Control | Spatial Pearson $r$ | Sign agreement |
| --- | --- | --- | --- |
| Visual features | Gabor | $0.9492 \pm 0.0041$ | $0.9257 \pm 0.0094$ |
| Visual features | GIST | $0.9556 \pm 0.0038$ | $0.9270 \pm 0.0103$ |
| Visual features | Sketch-token | $0.9628 \pm 0.0029$ | $0.9838 \pm 0.0016$ |
| Visual features | Combined | $0.9413 \pm 0.0038$ | $0.9147 \pm 0.0120$ |
| Object layout | Primary-object area | $0.9989 \pm 0.0001$ | $0.9965 \pm 0.0006$ |
| Object layout | Object count | $0.9999 \pm 0.0000$ | $0.9990 \pm 0.0003$ |
| Object layout | Non-primary-object area | $0.9957 \pm 0.0004$ | $0.9941 \pm 0.0019$ |
| Object layout | Joint layout | $0.9873 \pm 0.0005$ | $0.9925 \pm 0.0011$ |

**Table S2:** Incremental held-out variance explained by real-world size. Values report the across-subject mean increase in held-out explained variance obtained by adding real-world size to the combined visual-feature model. Statistical significance was assessed using two-sided one-sample *t*-tests across the eight subjects, followed by Benjamini–Hochberg FDR correction across the 13 ROIs. Note that these values reflect an average over all voxels per ROI (including voxels without size sensitivity), which leads to small magnitude of the average *R*^2^ values.

| ROI | Mean $\Delta R^2$ | SEM | $t_7$ | $q$ |
| --- | --- | --- | --- | --- |
| V1 | 0.00036 | 0.00014 | 2.610 | 0.0378 |
| V2 | 0.00083 | 0.00021 | 3.869 | 0.0100 |
| V3 | 0.00089 | 0.00025 | 3.599 | 0.0114 |
| hV4 | 0.00139 | 0.00037 | 3.747 | 0.0104 |
| RSC | 0.00415 | 0.00068 | 6.100 | 0.0012 |
| PPA | 0.00281 | 0.00038 | 7.322 | 0.00069 |
| OPA | 0.00290 | 0.00023 | 12.456 | 0.000064 |
| EBA | 0.00251 | 0.00029 | 8.525 | 0.00039 |
| FBA | 0.00295 | 0.00065 | 4.554 | 0.0049 |
| OFA | 0.00140 | 0.00044 | 3.173 | 0.0185 |
| FFA | 0.00191 | 0.00032 | 5.983 | 0.0012 |
| OWFA | 0.00060 | 0.00025 | 2.353 | 0.0509 |
| VWFA | 0.00336 | 0.00055 | 6.140 | 0.0012 |

## References

Agrillo, C., Beran, M. J., & Parrish, A. E. (2019). Exploring the Jastrow illusion in humans (Homo sapiens), rhesus monkeys (Macaca mulatta), and capuchin monkeys (Sapajus apella). Perception, 48 (5), 367–385. 10.1177/0301006619838181

Allen, E. J., St-Yves, G., Wu, Y., Breedlove, J. L., Prince, J. S., Dowdle, L. T., Nau, M., Caron, B., Pestilli, F., Charest, I., Hutchinson, J. B., Naselaris, T., & Kay, K. (2022). A massive 7T fMRI dataset to bridge cognitive neuroscience and artificial intelligence. Nature Neuroscience, 25 (1), 116–126. 10.1038/s41593-021-00962-x

Andrews, T. J., Watson, D. M., Rice, G. E., & Hartley, T. (2015). Low-level properties of natural images predict topographic patterns of neural response in the ventral visual pathway. Journal of Vision, 15 (7), 3. 10.1167/15.7.3

Bar, M. (2004). Visual objects in context. Nature Reviews Neuroscience, 5 (8), 617–629.

Benjamini, Y., & Hochberg, Y. (1995). Controlling the false discovery rate: A practical and powerful approach to multiple testing. Journal of the Royal Statistical Society: Series B (Methodological), 57, 289–300. 10.1111/J.2517-6161.1995.TB02031.X

Benson, N. C., Jamison, K. W., Arcaro, M. J., Vu, A. T., Glasser, M. F., Coalson, T. S., Van Essen, D. C., Yacoub, E., Ugurbil, K., Winawer, J., & Kay, K. (2018). The Human Connectome Project 7 tesla retinotopy dataset: Description and population receptive field analysis. Journal of Vision, 18 (13), 23. 10.1167/18.13.23

Blumenthal, A., Stojanoski, B., Martin, C. B., Cusack, R., & Köhler, S. (2018). Animacy and real-world size shape object representations in the human medial temporal lobes. Human Brain Mapping, 39, 3779–3792. 10.1002/HBM.24212

Chang, N., Pyles, J. A., Marcus, A., Gupta, A., Tarr, M. J., & Aminoff, E. M. (2019). BOLD5000, a public fMRI dataset while viewing 5000 visual images. Scientific Data, 6 (1), 1–18.

Cohen, L., Dehaene, S., Naccache, L., Lehéricy, S., Dehaene-Lambertz, G., Hénaff, M.-A., & Michel, F. (2000). The visual word form area: Spatial and temporal characterization of an initial stage of reading in normal subjects and posterior split-brain patients. Brain, 123 (2), 291–307.

Doherty, M. J., Campbell, N. M., Tsuji, H., & Phillips, W. A. (2010). The Ebbinghaus illusion deceives adults but not young children. Developmental Science, 13 (5), 714–721.

Epstein, R., & Kanwisher, N. (1998). A cortical representation of the local visual environment. Nature, 392 (6676), 598–601.

Epstein, R. A., & Baker, C. I. (2019). Scene perception in the human brain. Annual Review of Vision Science, 5, 373–397.

Gallivan, J. P., Cavina-Pratesi, C., & Culham, J. C. (2009). Is that within reach? fMRI reveals that the human superior parieto-occipital cortex encodes objects reachable by the hand. Journal of Neuroscience, 29 (14), 4381–4391.

Gao, J. S., Huth, A. G., Lescroart, M. D., & Gallant, J. L. (2015). Pycortex: An interactive surface visualizer for fMRI. Frontiers in Neuroinformatics, 9, 23. 10.3389/fninf.2015.00023

Gibson, J. (1950). The perception of the visual world. Houghton Mifflin. https://psycnet.apa.org/record/1951-04286-000

Grèzes, J., Tucker, M., Armony, J., Ellis, R., & Passingham, R. E. (2003). Objects automatically potentiate action: An fMRI study of implicit processing. European Journal of Neuroscience, 17 (12), 2735–2740.

Hebart, M. N., Contier, O., Teichmann, L., Rockter, A. H., Zheng, C. Y., Kidder, A., Corriveau, A., Vaziri-Pashkam, M., & Baker, C. I. (2023). THINGS-data, a multimodal collection of large-scale datasets for investigating object representations in human brain and behavior. eLife, 12, e82580. 10.7554/eLife.82580

Henderson, M. M., Tarr, M. J., & Wehbe, L. (2023). Low-level tuning biases in higher visual cortex reflect the semantic informativeness of visual features. Journal of Vision, 23, 8–8. 10.1167/JOV.23.4.8

Huang, T., Song, Y., & Liu, J. (2022). Real-world size of objects serves as an axis of object space. Communications Biology, 5 (1), 1–12.

Jain, N., Wang, A., Henderson, M. M., Lin, R., Prince, J. S., Tarr, M. J., & Wehbe, L. (2023). Selectivity for food in human ventral visual cortex. Communications Biology, 6 (1), 175.

Jiang, M., Huang, S., Duan, J., & Zhao, Q. (2015). SALICON: Saliency in context. Proceedings of the IEEE conference on computer vision and pattern recognition, 1072–1080.

Josephs, E. L., & Konkle, T. (2020). Large-scale dissociations between views of objects, scenes, and reachable-scale environments in visual cortex. Proceedings of the National Academy of Sciences, 117 (47), 29354–29362.

Julian, J. B., Ryan, J., & Epstein, R. A. (2017). Coding of object size and object category in human visual cortex. Cerebral Cortex, 27 (6), 3095–3109.

Kanwisher, N., McDermott, J., & Chun, M. M. (1997). The fusiform face area: A module in human extrastriate cortex specialized for face perception. Journal of Neuroscience, 17 (11), 4302–4311.

Konkle, T., & Caramazza, A. (2013). Tripartite organization of the ventral stream by animacy and object size. Journal of Neuroscience, 33 (25), 10235–10242.

Konkle, T., & Oliva, A. (2011). Canonical visual size for real-world objects. Journal of Experimental Psychology: Human Perception and Performance, 37 (1), 23.

Konkle, T., & Oliva, A. (2012). A real-world size organization of object responses in occipitotemporal cortex. Neuron, 74 (6), 1114–1124.

Künnapas, T. M. (1955). An analysis of the vertical-horizontal illusion. Journal of Experimental Psychology, 49 (2), 134–140. 10.1037/h0045229

Lescroart, M. D., & Gallant, J. L. (2019). Human scene-selective areas represent 3D configurations of surfaces. Neuron, 101 (1), 178–192. 10.1016/j.neuron.2018.11.004

Lim, J. J., Zitnick, C. L., & Dolĺar, P. (2013). Sketch tokens: A learned mid-level representation for contour and object detection. Proceedings of the IEEE conference on computer vision and pattern recognition, 3158–3165.

Lin, T. Y., Maire, M., Belongie, S., Hays, J., Perona, P., Ramanan, D., Dolĺar, P., & Zitnick, C. L. (2014). Microsoft COCO: Common objects in context. Lecture Notes in Computer Science (including subseries Lecture Notes in Artificial Intelligence and Lecture Notes in Bioinformatics), 8693 LNCS(PART 5), 740–755. 10.1007/978-3-319-10602-148

Long, B., & Konkle, T. (2017). A familiar-size Stroop effect in the absence of basic-level recognition. Cognition, 168, 234–242.

Long, B., Konkle, T., Cohen, M. A., & Alvarez, G. A. (2016). Mid-level perceptual features distinguish objects of different real-world sizes. Journal of Experimental Psychology: General, 145 (1), 95.

Long, B., Yu, C.-P., & Konkle, T. (2018). Mid-level visual features underlie the high-level categorical organization of the ventral stream. Proceedings of the National Academy of Sciences, 115 (38), E9015–E9024.

Mahon, B. Z., Milleville, S. C., Negri, G. A. L., Rumiati, R. I., Caramazza, A., & Martin, A. (2007). Action-related properties shape object representations in the ventral stream. Neuron, 55 (3), 507–520. 10.1016/j.neuron.2007.07.011

McCandliss, B. D., Cohen, L., & Dehaene, S. (2003). The visual word form area: Expertise for reading in the fusiform gyrus. Trends in Cognitive Sciences, 7 (7), 293–299.

Murray, S. O., Boyaci, H., & Kersten, D. (2006). The representation of perceived angular size in human primary visual cortex. Nature Neuroscience, 9 (3), 429–434.

Naselaris, T., Kay, K. N., Nishimoto, S., & Gallant, J. L. (2011). Encoding and decoding in fMRI. NeuroImage, 56 (2), 400–410.

Naselaris, T., Stansbury, D. E., & Gallant, J. L. (2012). Cortical representation of animate and inanimate objects in complex natural scenes. Journal of Physiology-Paris, 106 (5-6), 239–249. 10.1016/j.jphysparis.2012.02.001

Nasr, S., Echavarria, C. E., & Tootell, R. B. (2014). Thinking outside the box: Rectilinear shapes selectively activate scene-selective cortex. Journal of Neuroscience, 34 (20), 6721–6735. 10.1523/JNEUROSCI.4802-13.2014

Nasr, S., & Tootell, R. B. (2012). A cardinal orientation bias in scene-selective visual cortex. Journal of Neuroscience, 32 (43), 14921–14926.

Oliva, A., & Torralba, A. (2001). Modeling the shape of the scene: A holistic representation of the spatial envelope. International Journal of Computer Vision, 42 (3), 145–175.

Prince, J. S., Charest, I., Kurzawski, J. W., Pyles, J. A., Tarr, M. J., & Kay, K. N. (2022). Improving the accuracy of single-trial fMRI response estimates using GLMsingle (P. Kok, F. P. de Lange, P. Kok, & B. Turner, Eds.). eLife, 11, e77599.

Reddy, N., Jain, S., Yarlagadda, P., & Gandhi, V. (2020). Tidying deep saliency prediction architectures. 2020 *IEEE/RSJ International Conference on Intelligent Robots and Systems (IROS)*, 10241–10247.

Russakovsky, O., Deng, J., Su, H., Krause, J., Satheesh, S., Ma, S., Huang, Z., Karpathy, A., Khosla, A., Bernstein, M., Berg, A. C., & Fei-Fei, L. (2015). ImageNet large scale visual recognition challenge. International Journal of Computer Vision, 115 (3), 211–252. 10.1007/s11263-015-0816-y

Sergent, J., Ohta, S., & MacDonald, B. (1992). Functional neuroanatomy of face and object processing: A positron emission tomography study. Brain, 115, 15–36.

Slater, A., Mattock, A., & Brown, E. (1990). Size constancy at birth: Newborn infants’ responses to retinal and real size. Journal of Experimental Child Psychology, 49 (2), 314–322.

Sperandio, I., & Chouinard, P. A. (2015). The mechanisms of size constancy. Multisensory Research, 28 (3-4), 253–283. 10.1163/22134808-00002483

Stoinski, L. M., Konkle, T., & Hebart, M. N. (2026). Principles of coarse-scale functional organization in occipitotemporal cortex [Preprint]. bioRxiv. 10.64898/2026.02.18.706551

St-Yves, G., & Naselaris, T. (2018). The feature-weighted receptive field: An interpretable encoding model for complex feature spaces. NeuroImage, 180, 188–202. 10.1016/j.neuroimage.2017.06.035

Surkys, T., Bertulis, A., & Bulatov, A. (2006). Delboeuf illusion study. Medicina (Kaunas), 42 (8), 673–681.

Tucker, M., & Ellis, R. (2001). The potentiation of grasp types during visual object categorization. Visual Cognition, 8 (6), 769–800.

Warrington, E. K., & Shallice, T. (1984). Category specific semantic impairments. Brain, 107 (3), 829–853.

Xiao, J., Hays, J., Ehinger, K. A., Oliva, A., & Torralba, A. (2010). SUN database: Large-scale scene recognition from abbey to zoo. 2010 IEEE Computer Society Conference on Computer Vision and Pattern Recognition, 3485–3492. 10.1109/CVPR.2010.5539970

